# Perceived Control Over Breathing Perturbations Depends on Interosensory Input and Prediction Error

**DOI:** 10.64898/2026.07.30.741475

**Authors:** Alexander J. Hess, Giulia Cornali, Stephanie Mellor, László Demkó, Laura Köchli, Chiara Bassi, Alexandra Kalberer, Stephanie Marino, Christoph Mathys, Stefan Frässle, Jakob Siemerkus, Olivia K. Harrison, Jakob Heinzle, Sandra Iglesias, Klaas Enno Stephan

## Abstract

Interoception and homeostatic/allostatic control are not only fundamental for survival but play a key role for maintaining somatic and mental health. Furthermore, metacognitive evaluations of these processes, such as perceived failure of allostatic regulation, have been proposed to underlie the experience of chronic symptoms, such as chronic fatigue. A central question in this regard is what mechanisms the brain uses to evaluate control over bodily states. A prominent computational proposal posits that this is achieved by monitoring of interoceptive prediction errors (PEs). However, to date, this proposal has not been tested experimentally.

Here, we tested this hypothesis by applying computational process models of perceived control to data from a novel behavioural paradigm, the Respiratory Metacognition of Control Task (RMCT). The RMCT manipulates control over breathing by changing inspiratory resistive loads as a function of control achieved in a gamified prediction task. We developed and compared trial-by-trial generative models of perceived explicit control in the RMCT, using data from 50 volunteers in a pre-registered analysis. Bayesian model selection suggested that perceived control over breathing is best explained as a function of both trial-wise interoceptive outcomes (breathing with or without an inspiratory resistive load) and trial-wise PEs about respiratory resistance.

These results support a longstanding computational proposal of how the brain detects failures of bodily regulation and provide a mechanistic model for understanding inter-individual differences in perceived control over bodily states.

## Introduction

A number of constructs of control have been linked to mental health in the literature, with prominent theories, including Rotter’s locus of control (Rotter, 1966; Rotter and Mulry, 1965) and Bandura’s self-efficacy theory (Bandura, 1988, 1977). These frameworks have advanced our understanding of how individuals perceive their influence over life events and personal outcomes, and how these perceptions affect psychological well-being (Infurna et al., 2016; Luszczynska et al., 2005). In essence, perceived lack of control over oneself and one’s environment is associated with reduced well-being and is considered a risk factor for various mental health problems, including anxiety, depression, and suicidal ideation (for meta-analyses, see Cheng et al., 2013; Gallagher et al., 2014; Hirschmiller et al., 2024).

Recent developments in mental health research have shifted the focus from perception of the external world to the body’s internal environment (Heim et al., 2023; Khalsa et al., 2018; Nord and Garfinkel, 2022; Quadt et al., 2018; Tsakiris and Critchley, 2016). In parallel, perceived control over bodily states has become a central component in general frameworks of mental health that refer to computational (algorithmic) principles of brain-body interactions (Petzschner et al., 2017). A concrete case is the allostatic self-efficacy theory (ASE; Manjaly et al., 2019; Stephan et al., 2016) which conceptualises chronic fatigue as the brain’s metacognitive diagnosis that it lacks control over bodily states (low allostatic self-efficacy). More concretely (and slightly simplified), the ASE theory proposes that (i) fatigue arises when the brain detects a state of dyshomeostasis whose resolution requires the organism to rest, (ii) the detection of dyshomeostasis relies on monitoring a single computational quantity, interoceptive Shannon surprise or, equivalently, interoceptive prediction errors (PEs), and (iii) fatigue becomes chronic when no regulatory action, including rest, succeeds in reducing interoceptive PEs, entrenching the feeling of lack of control over bodily states (Stephan et al., 2016). While empirical evidence supporting several central predictions of the ASE theory has been provided (Hess et al., 2024; Rouault et al., 2023), one prediction has remained untested: that the experience of control over bodily states is a function of interoceptive PEs.

Given the general importance of perceived control for mental health (see above), testing this prediction is of general interest, over and beyond the ASE theory. So far, empirical investigations have been difficult due to the lack of both suitable experimental paradigms and process models that describe how an individual’s perception of control over bodily states arises. In this study, we address both issues, presenting both a paradigm and a process model.

Specifically, capitalising on previous developments for investigating interoceptive PEs in the realm of breathing (Harrison et al., 2021), we developed the Respiratory Metacognition of Control Task (RMCT). The RMCT is a behavioural paradigm for manipulating one’s control over respiratory resistance while recording both trial-wise predictions and experience of control. Additionally, we propose a process model for perceived control over bodily (respiratory) states that considers both interoceptive PEs and trial-wise outcomes. In pre-registered analyses, we compared several variations of our model and a control model, using data acquired from a total of 50 healthy individuals.

## Methods

In the following section, we describe (i) the experimental setup and the study population, and (ii) the computational model and statistical data analysis. Notably, we prespecified the hypotheses and analyses methods in a pre-registered analysis plan (available at https://doi.org/10.5281/zenodo.15388133) prior to data analysis. Please note that, for consistency, we reused text from our analysis plan in this manuscript, in adapted and extended form.

### Research Questions

Our primary research question was whether trial-by-trial perceived control over breathing can be explained as a function of interoceptive prediction errors (PEs) in the RMCT. This was complemented by a set of secondary research questions, i.e. whether task-based readouts from the RMCT – which induces transient experiences of lack of control – are related to more stable constructs of fatigue and perceived control measured by questionnaires. These secondary questions were exploratory since it is unclear whether and to what degree transient (state-like) aspects of perceived control are related to stable (trait-like) aspects of mental health.

Concretely, our research questions (as pre-registered in the analysis plan) were:

1. Can perceived control over breathing be explained as a function of interoceptive prediction errors in the RMCT?
2. Is the experience of control during the RMCT related to the overall experience of fatigue?
3. Is the experience of control during the RMCT related to physical, cognitive, and psychosocial dimensions of fatigue?
4. Can the explanation of fatigue scores via sleep and interoception (see Rouault et al., 2023) be improved by taking into account the experience of control during the RMCT?
5. Is the experience of control during the RMCT related to other constructs of control?

### Study Design and Population

The study was approved by the Ethics Committee of the Canton of Zurich (BASEC number: 2022-02178). We acquired behavioural, physiological and questionnaire data from a total of N=50 healthy volunteers. The data were split into a discovery set and a validation set. The discovery set included *N*_ds_=20 volunteers (12 females, 8 males; age 25.2±4.9 (mean + standard deviation) years) for data exploration and the estimation of empirical priors. The validation set contained data from *N*_vs_=30 volunteers (18 females, 12 males; age 23.7 ± 4.1 years). The sample size was determined using a Sequential Bayes Factor (SBF) design (Schönbrodt et al., 2017) based on our primary hypothesis test, as outlined in the statistical analysis plan. Discovery and validation data sets remained strictly separated to prevent any information leakage.

For our secondary questions, we aimed for a study population with different levels of fatigue and a balanced gender ratio. Participants were therefore pre-screened with the Fatigue Assessment Scale (FAS; Michielsen et al., 2004) and selected in order to maximise variance of fatigue scores (discovery set: FAS 22.4 ± 6.9, validation set: FAS 21.8 ± 5.8) while keeping a balanced gender ratio. Upon study inclusion and before coming to the lab, participants answered demographic as well as socioeconomic questions and completed the following battery of psychological online questionnaires covering depressive symptoms (Patient Health Questionnaire, PHQ-8; Löwe et al., 2002), trait anxiety (Spielberger State-Trait Anxiety Inventory, STAI-T; Laux, 1981), general self-efficacy (General Self-Efficacy Scale, GSES; Schwarzer, 1993; Schwarzer et al., 1997; Schwarzer and Jerusalem, 1999, 1995), interoception (Multidimensional Assessment of Interoceptive Awareness Version 2, MAIA-2; Mehling et al., 2018, 2012), control (Perceived Control Items, PCI; Infurna et al., 2011), locus of control (LOC; Krampen, 1985, 1981, 1979), and sleep (Pittsburgh Sleep Quality Index, PSQI; Buysse et al., 1989; Hinz et al., 2017). On the day of the experiment, participants filled in two questionnaires assessing fatigue (FAS; Michielsen et al., 2004; Modified Fatigue Impact Scale, MFIS; Zimmermann and Hohlfeld, 1999) and completed the RMCT followed by a number of task-specific debriefing questions.

### The Respiratory Metacognition of Control Task (RMCT)

Figure 1A displays a graphical summary of the RMCT. Participants were asked to navigate an avatar jumping from one island to the next by selecting the strength of the jump at a fixed jump angle of 45 degrees. The task consisted of 80 trials in total, with varying distance between the two islands and the target island additionally varying in size (large/small), creating different levels of task difficulty. In 47.5% of the trials, a small wind symbol indicated the presence of wind during that trial, leading to a deviation of the jump trajectory in either forward or backward direction. The direction of the wind was unknown to the participant while selecting the jump strength, thus adding uncertainty about the avatar’s landing position. All participants were presented with identical stimuli (island distance, island size, presence/absence of wind) in the same order; see colour-coding in Figure 1B for the trajectory of task conditions. Using a fixed task trajectory is a typical choice in computational studies of learning tasks as it ensures that parameter estimates refer to the same type of learning process and can be compared across participants.

**Figure 1.**
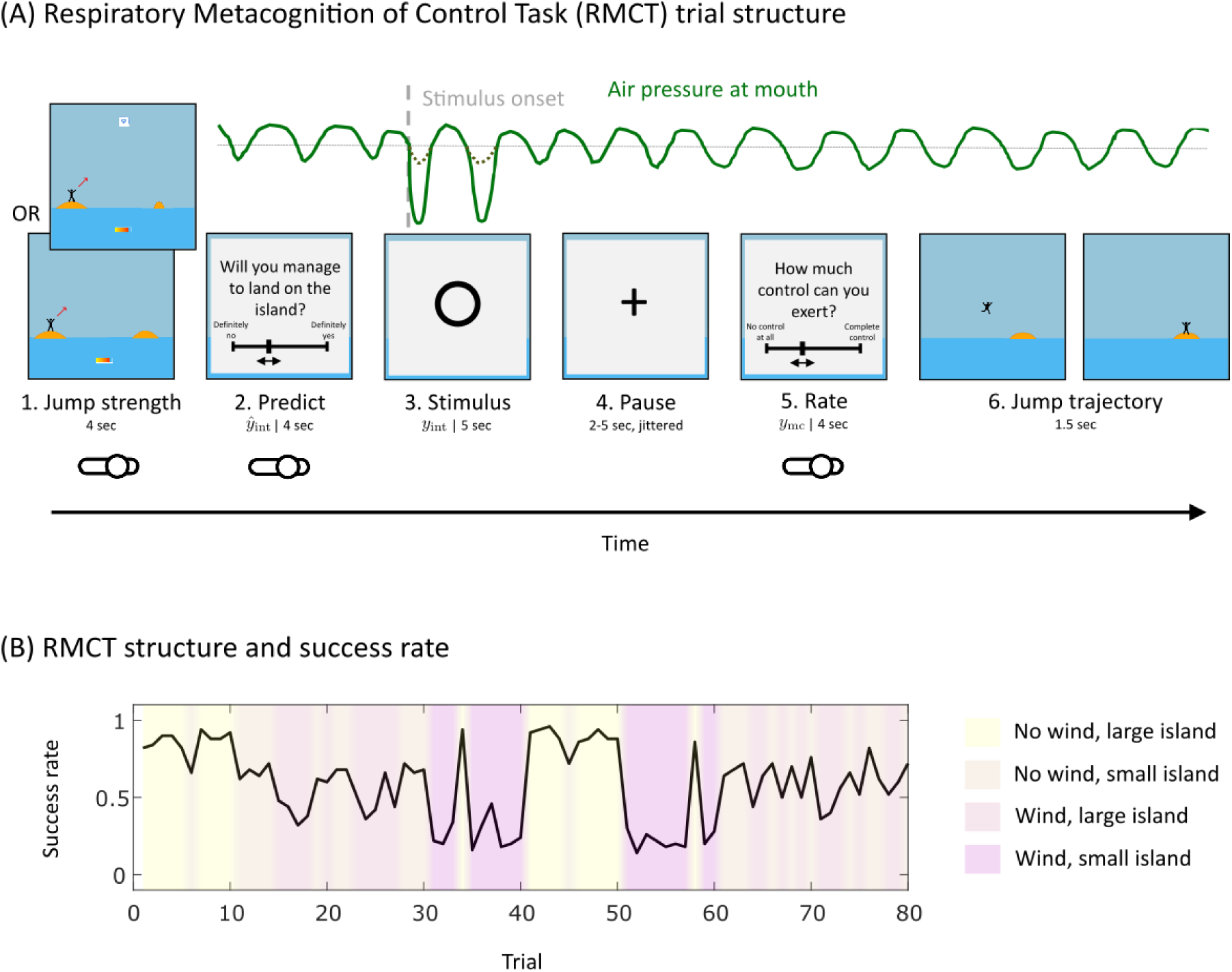
Task structure of the Respiratory Metacognition of Control Task (RMCT). (A) Overview of the structure of one trial of the RMCT. The questions displayed (2. Predict and 5. Rate) are only placeholders, for exact wording, please refer to the main text. (B) displays the success rate across all participants (*N*_tot_ = 50) for each of the 80 trials, colour-coded by trial condition.

During the entire task, participants were lying supine with a monitor positioned vertically above their head. They wore a 7450 Series Silicone V2™ Oro-Nasal Mask (Hans Rudolph Inc., Lenexa, USA) connected to a mechanical breathing system that allows for remote administration and monitoring of inspiratory resistive loads (technical details of the breathing circuit used for inspiratory resistive loading are described in Rieger et al., 2020 and a schematic overview of the breathing circuited is provided in Supplementary Figure S6).

The within-trial structure is shown by Figure 1A. On every trial, participants were asked to specify the jump strength required to land on the target island using a mechanical slider with their right hand. Next, a pop-up window appeared on the screen asking the participants to predict the outcome of a given trial: “Are you able to land on the island and thereby prevent a breathing resistance?”, on a visual analogue scale (VAS) from “definitely no” to “definitely yes”. We refer to this prediction as *y*^_int_. Following this prediction, a circle appeared on the screen to indicate the interoceptive stimulus period (5s). During this period, participants either experienced a positive outcome (*y*_int_ = 1), where no added resistive load was applied if they successfully landed on the island, or a negative outcome (*y*_int_ = 0) where they had to breathe against an added inspiratory resistive load if they missed the target island.

The inspiratory resistive load was administered at 70% of each subject’s individual maximal inspiratory pressure, as measured in the laboratory, and was delivered via a PowerBreathe KH2 (PowerBreathe International Ltd, Warwickshire, UK). This ensured that participants perceived a relevant increase in inspiratory resistance. Our setup for remote automated inspiratory resistive loading was based on the breathing circuit developed by Rieger et al. (2020) and used by previous studies (Brand et al., 2024; Harrison et al., 2021). We extended the previously used breathing circuit such that the onset of the 5s breathing stimulus period was coupled to the onset of inspiration in the breathing cycle, to ensure that participants perceived an increase in inspiratory resistance clearly. This was accomplished by using the measured air flow (output from a spirometer connected to a respiratory flow head 300 l; ADInstruments Limited, Oxford, United Kingdom) as an input to a custom-built algorithm for real-time control on an Arduino Uno R3 microcontroller (https://www.arduino.cc/). Following the period of breathing stimulation, participants were given a variable pause of 2–5s before being asked to rate their perceived control over breathing perturbation by answering “How well can you actively control whether you receive a breathing resistance?” on a VAS scale from “no control at all” to “complete control”. We refer to this subjective rating of perceived control over breathing resistances as *y*_mc_. Afterwards, the pop-up window closed and participants were presented with the full jump trajectory and landing position before moving on to the next trial. This enabled trial-by-trial reflection about task performance and improvement thereof.

Before conducting the experiment, participants were presented with standardised instructions. For familiarisation with the task, we employed 20 practice trials (without the mechanical breathing system) and an additional 6 trials for familiarisation with the inspiratory resistive loadings. Task and instructions were both presented in German (in this manuscript, present the English translation. The original German wordings can be found in Supplementary section S7).

After every 10^th^ trial of the task, participants were additionally asked: “Did you feel in control over your breathing during the previous trials?” on a VAS from “definitely no” to “definitely yes”, and “How easy was it for you to tolerate the resisted breathing during the previous trials?” on a VAS from “not easy at all” to “very easy”. We refer to these ratings as *y*_c_ (overall control over breathing) and *y*_tol_ (tolerance of resisted breathing), respectively. Immediately after the last trial of the task, participants were asked to rate ‘‘How unpleasant did you find the breathing resistances during the task?’’ on a VAS from ‘‘not at all unpleasant” to ‘‘maximally unpleasant”. We refer to this unpleasantness rating as *y*_av_.

During the entire task, we recorded the following physiological signals: pressure at the mouth, respiratory gas concentrations (O2 and CO2), air flow and pulse. These physiological signals are not of relevance for the hypotheses tested in the current study and their analysis is therefore beyond the scope of the current manuscript.

### Computational Modelling of Behaviour

We developed a set of trial-by-trial computational models of behaviour representing competing hypotheses. Our pre-registered analyses followed a Bayesian workflow for generative modelling of behavioural data, as described previously (Hess et al., 2025) including specification of an initial model space, the choice of suitable prior configurations, the choice of model inversion technique and its validation, model inversion given the empirical data set, model comparison (hypothesis testing), and model evaluation.

#### Model space

Our primary research question (RQ1) was whether the experience of control over inspiratory resistive loads could be explained as a function of interoceptive prediction errors. We formalised this in the form of a generative model for trial-by-trial reports of perceived control over breathing. We reasoned that the experience of control would not occur (or disappear) instantaneously when interoceptive PEs were present or absent but represented a time-weighted average of past experiences of PEs. We further reasoned that, in addition to PEs, the actual interosensory input (i.e. whether there was respiratory resistance or not, independent of predictions) would contribute to the experience of control, in a time-weighted fashion as for PEs.

A generic approach to formulate the notion of time-weighting is an AR(1) model (autoregressive model of order 1) where the influence of past experience decays exponentially in time. Written recursively, our model is:

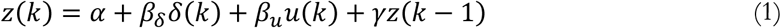

Here, *z* represents the participant’s expressed rating of perceived control over breathing perturbation on each trial *k*. *α* is a constant offset, *β_δ_* is a weighting factor for the influence of the prediction error 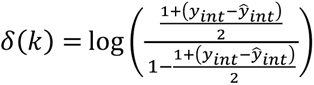 describing the difference between observed interosensory input *y*_int_ and predicted interosensory input *y*^_int_ at trial *k*, transformed to logit space. *β_u_* weighs the influence of the interosensory input *u* = 2*y*_int_ − 1 at trial *k*, recoded to {−1,1} corresponding to bad/good outcomes or the presence/absence of an inspiratory resistive load, respectively. γ represents the dependence on the experience of control over breathing perturbation from the previous trial *k* − 1. Equation 1 can be rewritten in an explicit fashion by solving the recursive formula and subsequent reparameterisation (for the full derivation, refer to the statistical analysis plan):

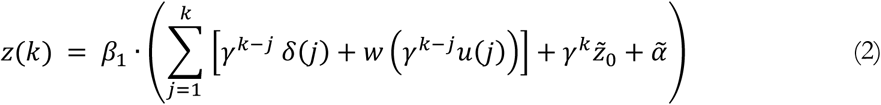

with 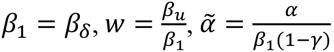, and 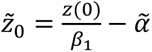.

The parameters in the reparameterised model (Eq. 2) can be interpreted as follows: *z̃*_0_ corresponds to the rescaled and shifted initial value of perceived control over breathing perturbation at trial zero. γ represents the dependence of the current experience of control on past experience of control. The latter, in turn, depends on past values of *δ* and *u*, and on *z̃*_0_. For this reason, γ in Eq. 2 denotes the influence of multiple factors from the past. *w* defines the influence of the weighted history of interosensory inputs *u* obtained during the task. *β*_1_ represents the sensitivity to *δ* and *u*, and *⍺̃* represents the rescaled tendency to experience high or low control.

To answer our primary research question, we compared the full model (***M****_δ_*_,*u*_ in Table 1), which included both *δ* and *u* as predictor variables from Eq. 2, to a null model (***M***_0_) which assumed perceived control to be constant over time within a given individual but allowed it to differ across individuals. Additionally, we also considered reduced versions of the full model ***M****_δ_*_,*u*_ which contained only one of the two predictors (***M****_u_*: *u*; ***M****_δ_*: *δ*). Table 1 lists the exact equations of all four models.

**Table 1.** Equations of trial-wise ratings of perceived control over breathing perturbation *z*(*k*) for all four models in the model space. *k* indexes the current trial, *α* is a constant offset, *β*_1_ represents the sensitivity to the logit-transformed prediction error *δ* and the recoded interosensory input *u*. and *⍺̃* represents the rescaled tendency to experience high or low control. *z̃*_0_corresponds to the rescaled and shifted initial value of perceived control over breathing perturbation at trial zero and *⍺̃* represents the rescaled tendency to experience high or low control. γ in denotes the influence of multiple factors from the past. *w* defines the influence of the weighted history of interosensory inputs *u* obtained during the task.

| Model | Perceived control over breathing perturbation $z$ |
| --- | --- |
| $M_0$ | $z(k) = \alpha$ |
| $M_u$ | $z(k) = \beta_1 \cdot (\sum_{j=1}^k \gamma^{k-j} u(j) + \gamma^k \tilde{z}_0 + \tilde{\alpha}), \quad \text{with } \beta_1 = \beta_u$ |
| $M_\delta$ | $z(k) = \beta_1 \cdot (\sum_{j=1}^k \gamma^{k-j} \delta(j) + \gamma^k \tilde{z}_0 + \tilde{\alpha}), \quad \text{with } \beta_1 = \beta_\delta$ |
| $M_{\delta,u}$ | $z(k) = \beta_1 \cdot \left( \sum_{j=1}^k [\gamma^{k-j} \delta(j) + w(\gamma^{k-j} u(j))] + \gamma^k \tilde{z}_0 + \tilde{\alpha} \right)$ |

Under the assumption of normality of the trial-wise logit-transformed ratings of perceived control over breathing resistances *y*_mc_, the trial-wise likelihood function of the generative model was defined as follows:

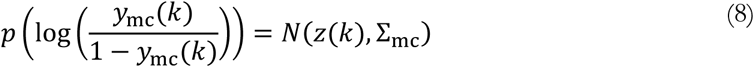

where 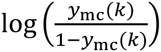 corresponds to a logit-transformation of the perceived control over breathing perturbation rating *y*_mc_ at trial *k*. To get a full generative model, we defined a prior probability distribution over the parameter space **θ** = (*γ*, *w*, *z̃*_0_, *β*_1_, *⍺̃*, Σ*_mc_*). We assumed that the prior distribution factorises into marginal priors

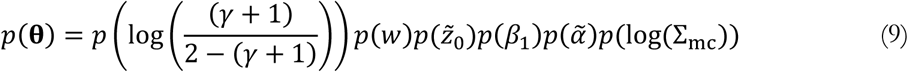

where each marginal prior corresponds to a normal distribution.

#### Prior elicitation

The sufficient statistics of the marginal *initial prior* densities are listed in our statistical analysis plan. These initial priors were used for inversion of the generative model for the discovery set (*N*_ds_=20) and were chosen to be uninformative to avoid overly strong assumptions. The estimated posterior means were then used as the sufficient statistics of *empirical prior* densities as the basis for model inversion using the validation data set (Table 2). This established two-step procedure enabled the use of informative priors covering a plausible range of human behaviour while avoiding the problem of double-dipping (Harrison et al., 2021; Hess et al., 2025; Schöbi et al., 2021). Visualisations of *initial prior* densities, maximum a posteriori (MAP) estimates from model inversion on the discovery set under the *initial priors*, as well as estimated *empirical prior* densities are provided in the Supplementary Materials (section S3).

**Table 2.** Sufficient statistics of the marginal *empirical prior* densities. All parameters have Gaussian marginal priors. *Empirical prior* means (***pE***) and variances (***pC*)** are listed for each model (rows) and parameter (column).

|  |  | parameters |  |  |  |  |  |
| --- | --- | --- | --- | --- | --- | --- | --- |
| | | $\log\left(\frac{(\gamma + 1)}{2 - (\gamma + 1)}\right)$ | $\tilde{\alpha}$ | $\beta_1$ | $\tilde{z}_0$ | $w$ | $\log(\Sigma_{mc})$ |
| $\mathbf{M}_0$ | $\mathbf{pE}_0$ | - | 0.34 | - | - | - | 0.28 |
| | $\mathbf{pC}_0$ | - | 0.73 | - | - | - | 0.38 |
| $\mathbf{M}_u$ | $\mathbf{pE}_u$ | 1.03 | 0.38 | 0.49 | -0.04 | - | -0.13 |
| | $\mathbf{pC}_u$ | 1.32 | 3.23 | 0.06 | 0 | - | 0.46 |
| $\mathbf{M}_\delta$ | $\mathbf{pE}_\delta$ | 0.72 | 1.51 | 0.05 | -0.04 | - | 0.30 |
| | $\mathbf{pC}_\delta$ | 2.71 | 10.72 | 0.05 | 0 | - | 0.41 |
| $\mathbf{M}_{\delta,u}$ | $\mathbf{pE}_{\delta,u}$ | 0.34 | -0.26 | -0.92 | -0.04 | -1.56 | -0.82 |
| | $\mathbf{pC}_{\delta,u}$ | 0.35 | 0.43 | 0.29 | 0 | 0.07 | 0.64 |

#### Model inversion and validation of computation

We inverted the generative models of behaviour using MAP estimation. The covariance of the approximate posterior was obtained under a Laplace approximation to the negative log joint at the MAP (Daw, 2011) as implemented in the HGF Toolbox (Frässle et al., 2021; Mathys et al., 2011, 2014). This allowed for calculation of an approximate log model evidence (LME) as basis for model comparison. To prevent the optimisation algorithm from getting stuck in local extrema of the objective function, we used a multi-start optimisation with 200 different starting points for the inversion of each subject and model, sampled from the respective *empirical prior* densities.

We validated our inference algorithm by assessing practical identifiability both at the level of parameters and models, using recovery analyses (Hess et al., 2025; Wilson and Collins, 2019). For this, we generated a synthetic data set (*N*_sim_=100) by randomly sampling parameter values from the *empirical prior* densities and plugging these values into the likelihood function. We inverted our models using the generated data under the respective *empirical priors*. Parameter recovery was assessed both qualitatively and quantitatively (Pearson correlation coefficients *ρ*) by comparing original (i.e. data-generating) parameter values to MAP estimates obtained using the data-generating model. Model identifiability was quantified both as the proportion of correctly identified models according to approximate LME scores in a classification analysis, and calculating protected exceedance probabilities (PXP) as part of random-effects (RFX) Bayesian model selection (BMS) (Rigoux et al., 2014; Stephan et al., 2009). Moreover, we computed log Bayes Factors (BFs) comparing the full model (***M****_δ_*_,*u*_) and the null model (***M***_0_) for synthetic data generated by either model.

#### Testing our primary hypothesis

To test our primary hypothesis that perceived control over breathing can be explained as a function of interoceptive PEs in the RMCT, we performed a fixed effects Bayesian Model Selection (FFX BMS) analysis. (FFX BMS, as opposed to random effects BMS, was chosen as primary approach since the Sequential Bayes Factor design of our study required that a single Bayes factor could be computed for the entire group.) We computed Group Bayes Factors (GBFs) between the full model ***M****_δ_*_,*u*_ and the null model ***M***_0_ for the validation set (*N*_vs_=30). For the interpretation of Bayes Factors, we used the grades of evidence according to the proposed scale by Kass and Raftery (1995), as specified in the analysis plan. Since FFX BMS is known to be sensitive to outliers (Stephan et al., 2009), we visualised the distribution of individual log BFs for the validation set. Additionally, we assessed the relative importance of the different components of ***M****_δ_*_,*u*_ as well as the prevalence of individual models in the population using RFX BMS.

#### Model evaluation

We qualitatively assessed individual-level and average model fits via visual inspection of the trajectories. Visualisation of the distribution of the residuals served as additional sanity check. We chose not to pre-specify any model evaluation procedures as we were uncertain a priori as to which metrics would prove the most informative. After observing the results from simulation analyses as well as our primary hypothesis test, we performed a number of analysis steps to improve our understanding of the obtained results and improve our confidence therein. To this end, we examined MAP estimates as well as contributions of the different components in our autoregressive models for perceived control over breathing perturbation. We compared the sign of the weighting terms between different models to obtain a potential explanation for the differences in goodness of fit that we observed.

### Testing our secondary hypotheses

Our additional hypotheses concerned the question of whether transient state-like readouts from the RMCT are related to more stable, trait-like constructs of fatigue (Hypotheses 2–4) and perceived control (Hypothesis 5) as measured by established questionnaires. We performed a set of Bayesian Linear Regression (BLR) analyses for different dependent variables (questionnaire scores) of the validation set (*N*_vs_=30). The independent variables consisted of a subset of variables from the RMCT that were pre-selected using the discovery set (*N*_ds_=20; see our analysis plan for a detailed description of the pre-selection procedure). The analyses of Hypotheses 2–5 all followed identical structure but for different dependent variables. For the sake of brevity, we provide a detailed description of this analysis procedure for one example, namely the analysis of Hypothesis 5 (H5). For a detailed overview of H2-H5, please refer to our statistical analysis plan.

In H5, we proposed that mean **ȳ***_c_* and variances *Var*(**y*_c_***) of overall control ratings from the RMCT explained the locus of control (LOC) questionnaire subscale related to the influence of ‘powerful others’ (LOC_p). The LOC_p score was therefore selected as the dependent variable in the BLR. We prespecified the Bayes factor *BF*_10_ as the test statistic of interest, comparing the full model *M*_1,H5_ containing the entire subset of predictors (**ȳ***_c_*, *Var*(**y*_c_***), gender, age, intercept) against a reduced model *M*_0,*H*5_ containing only gender, age, and an intercept. As exploratory analyses, we computed Bayes factors for all nested models of *M*_1,H5_ against *M*_0,H5_. We conducted an additional analysis using the score of the Perceived Control Items (PCI) questionnaire instead of the LOC_p score as dependent variable together with the respective subset of predictors identified on the discovery set.

### Deviations from the pre-registered analysis plan

In the process of our internal code review upon completion of the analyses, we discovered the following two errors: An error in the scoring of the PSQI questionnaire as well as an indexing error in the analysis of the questionnaire data leading to participants being associated with incorrect data. These issues affected the tests of our secondary hypotheses. As a consequence, we recomputed the entire analysis including the selection of independent variables on the discovery set, which resulted in a different subset of predictors for secondary hypotheses H2-H5, compared to those listed in the analysis plan. The results from the corrected version of the analysis and the updated set of predictors for the different BLRs are reported in Table 3. For those dependent variables where the re-analysis yielded no relevant predictors on the discovery set, we did not conduct any tests of the validation set and consequently did not list them in Table 3.

**Table 3.** Results of the analyses for hypotheses 4 and 5. Displayed are the dependent variables for each Bayesian Linear Regression (BLR) of Hypotheses 4 and 5 (Hyp *i*). *BF*_10_ denotes the Bayes Factor comparing *M*_1,*Hi*_ (covariates additional to *M*_0,*Hi*_ listed in the respective column) and *M*_0,*Hi*_ (covariates and factors additional to an intercept term listed in the respective column. Hypotheses 2 and 3 are not listed here because we did not find any relevant predictors in the discovery dataset and therefore did not conduct any tests for the validation dataset.

| Hyp $i$ | Dependent Variable | $M_{1,Hi}$ | $M_{0,Hi}$ | $BF_{10}$ |
| --- | --- | --- | --- | --- |
| $i = 4$ | MFIS | $\gamma$ | MAIA-2_38, PSQI, age, gender | 3.122 |
| $i = 5$ | PCI | $\bar{y}_c$ | age, gender | 0.584 |
| | LOC_p | $\bar{y}_c, Var(y_c)$ | age, gender | 1.483 |

Furthermore, the BLR analyses represent a deviation from our statistical analysis plan where we had planned to perform Bayesian ANCOVAs instead. However, we decided that BLR would be more appropriate to test our research questions at hand (please refer to section S5 in the Supplementary Materials for a detailed discussion of the matter.

## Data and Code Availability

The code implementing our statistical analysis is provided at https://gitlab.ethz.ch/tnu/code/hessetal_metac_analysis (will be made available upon acceptance for publication). The entire analysis pipeline was implemented in MATLAB R2024b (The MathWorks, Natick, MA, USA) and JASP (Version 0.19.1, https://jasp-stats.org/). Open-source software packages used for the analysis were added to the code repository as submodules referencing the utilised version. These packages include the HGF Toolbox (v7.1) as part of TAPAS (Frässle et al., 2021), the Variational Bayesian Analysis Toolbox (VBA, Daunizeau et al., 2014), and the DataViz library (https://doi.org/10.5281/zenodo.12749045). A researcher not involved in the initial data analysis performed an internal code review for quality assurance and to improve readability as well as usability of the code (Pereira et al., 2026). Study data were collected and managed using REDCap (Research Electronic Data Capture; Harris et al., 2019, 2009) electronic data capture tools hosted at ETH Zurich. The anonymised behavioural data set was published on Zenodo (https://doi.org/10.5281/zenodo.21700744; will be made available upon acceptance of the manuscript for publication) in a form adhering to the FAIR (Findable, Accessible, Interoperable, and Re-usable) data principles (Wilkinson et al., 2016). The task code is provided at https://gitlab.ethz.ch/demkol/metactask (will be made available upon acceptance for publication).

## Results

### Primary Hypothesis

The log group Bayes factor (GBF) for comparing the full model (***M****_δ_*_,*u*_) against the null model (***M***_0_) in the validation dataset was 1159, and the log average Bayes factor (the log of the geometric mean of the GBF) was 38.6. This result corresponds to ‘very strong’ evidence in favour of our primary hypothesis and illustrates that perceived explicit control over breathing can be explained as a function of interoceptive prediction errors (PEs), in combination with interoceptive inputs per se, in the RMCT. The distribution of individual log BFs in the validation dataset is shown in Figure 2A. This plot shows that the superiority of the full model was highly consistent across participants (for context, it is worth noting that a log BF of 3 or more is considered “very strong” evidence in favour of one model; Kass and Raftery, 1995).

**Figure 2.**
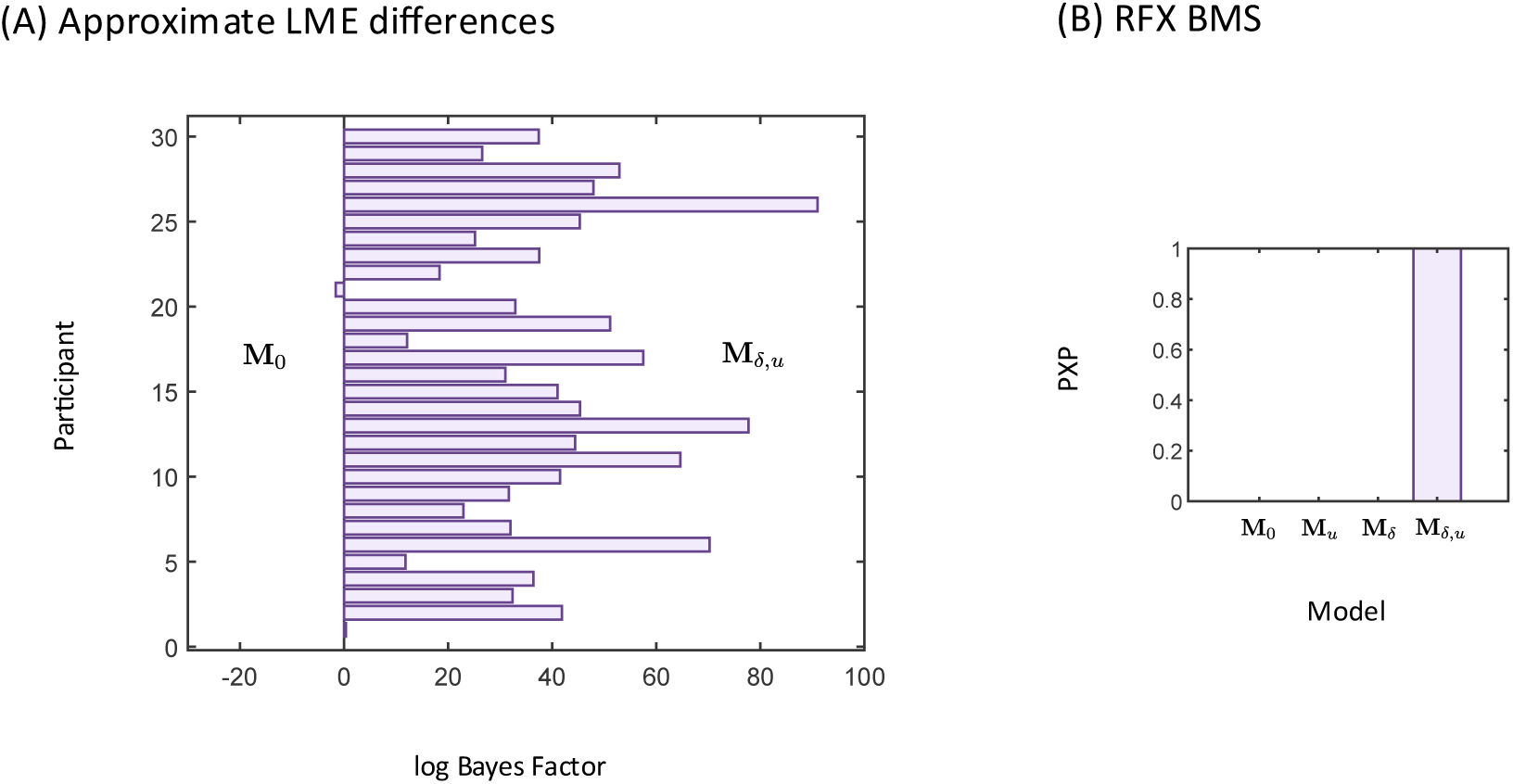
Primary hypothesis test. Log Bayes Factors (BFs) for comparing the full model (***M****_δ_*_,*u*_) and the null model (***M***_0_) for all 30 participants of the validation set are shown in (A). Protected exceedance probabilities (PXP) from RFX BMS applied to all four models are shown in panel (B).

In a subsequent step, we examined this result in more depth, using random effects Bayesian model selection (RFX BMS) to assess whether all components of the full model were needed to explain the behavioural data. To this end, we compared four models: the null model (***M***_0_), a model considering only interoceptive prediction errors (***M****_δ_*), a model considering only interoceptive inputs (***M****_u_*), and the full model which included both interoceptive prediction errors and inputs (***M****_δ_*_,*u*_). Comparing these four models against each other, the full model (***M****_δ_*_,*u*_) provided the best explanation for the dataset, showing a protected exceedance probability (PXP) of 1. The PXP denotes the probability that a particular model is more likely than all other models considered to have generated the data, corrected for the possibility that differences between models could have arisen by chance; see Rigoux et al. (2014) for details.

Visualisations of average model fits illustrated that ***M****_δ_*_,*u*_ provided an excellent fit to the participants’ expressed trial-wise ratings of perceived control; additionally, the histogram of residuals appeared to be approximately normally distributed (Figure 3). Additional plots of individual participant fits are shown in Figure 5A and in the Supplementary Material: To give an impression of the range of model fitting quality, Supplementary Figures S4A-B juxtapose the participants with the best and worst model fit (based on coefficient of determination). Quantitatively, model fit for individual participants (expressed as a Peasrson correlation coefficient and coefficient of determination) ranged from *ρ* = 0.41, *R*^2^ = 0.15 (worst model fit) to *ρ* = 0.97, *R*^2^ = 0.93 (best model fit) across our sample. The distribution of posterior means of model parameters is shown in Figure 4, both for the explicit and the recursive form of the model. As can be seen most easily in the plot for the recursive form of the model, on average, we found a positive effect of interosensory inputs *u* (where normal breathing=1 and restricted breathing=-1; see Methods) and a negative effect of signed PEs *δ* on ratings of perceived control. Validation analyses of our inference procedure revealed that the full model showed excellent parameter recoverability (*ρ* > 0.87) (see Supplementary materials, sections S1-S2 for details of parameter recovery and model identifiability analysis results).

**Figure 3.**
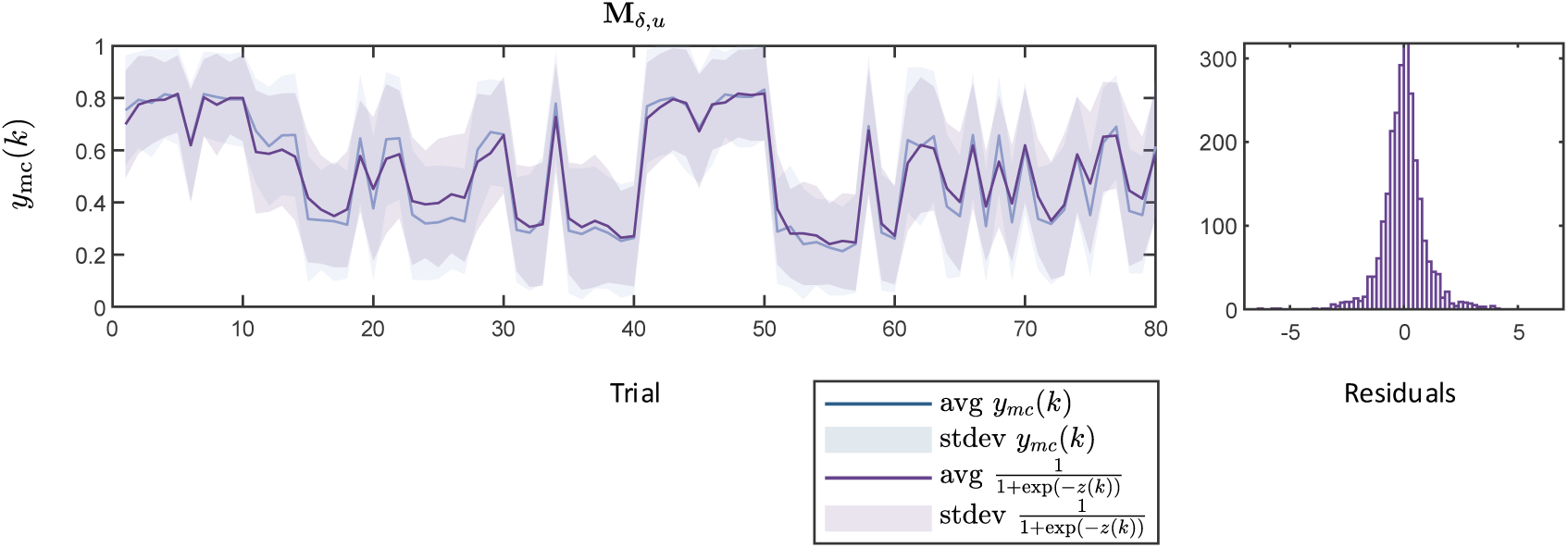
Group averages of data and model fit for the validation dataset (*N*_vs_ = 30). Left panel: Time series of average ratings of perceived control over breathing perturbations, *y*_mc_ (blue), and sigmoid transformed model predictions *z* (purple) for the full model ***M****_δ_*_,*u*_. Right panel: a histogram of model residuals (in logit-space).

**Figure 4.**
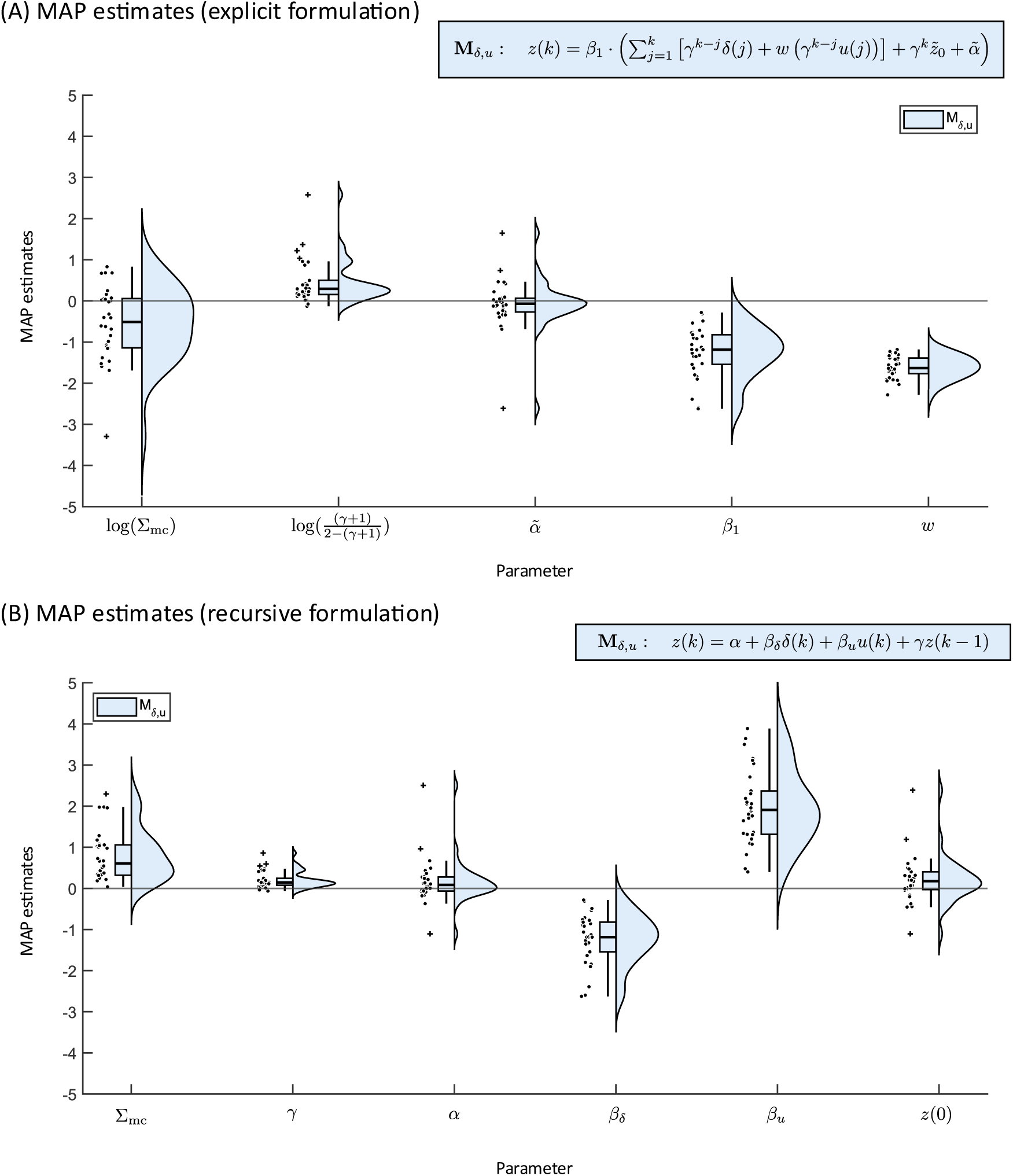
MAP estimates for the full model ***M****_δ_*_,*u*_ across validation set participants. (A) MAP estimates of all parameters in estimation space for the explicit model formulation (Eq. 2). (B) MAP estimates of selected parameters in native space for the recursive model formulations (Eq. 1).

### Additional Hypotheses

The pre-selection of regressors in the discovery set left us with three Bayesian linear regressions (BLRs) to be computed as part of the analyses for Hypotheses 2–5 for the dependent variables: MFIS, PCI, and LOC_p (powerful others). An overview of the results for H2-5 is provided in Table 3. We found ‘positive’ evidence (3 < *BF* ≤ 20) for Hypothesis 4 on the validation set, according to the interpretation of Bayes Factors provided by Kass and Raftery (1995). Here we had hypothesised that model-based estimates of perceived control over breathing perturbation, other ASE-inspired readouts (overall control, tolerance) and task unpleasantness in the RMCT would improve the explanation of fatigue scores from standard questionnaires (FAS, MFIS) when added to questionnaire scores (sleep and interoception; PSQI and MAIA-2, respectively) which have previously been used to explain fatigue scores (Rouault et al., 2023). The BLR for the dependent variable fatigue (measured by the MFIS) found ‘positive’ evidence (*BF*_10_ = 3.122) in favour of the model containing *γ* (the weighting parameter of past experience of control) as an additional predictor as opposed to the reduced model containing variables MAIA-2 (sum of subscales 3 and 8), PSQI, age and gender as well as an intercept. The full results of the BLR of H4 including details on the model comparison and posterior summaries of coefficients are reported in Supplementary Tables 2 and 3.

The analysis of Hypothesis 5 yielded ‘barely worth mentioning’ (i.e., inconclusive) levels (|*BF*| ≤ 3) of evidence either in favour of the best nested model tested (LOC_p) or in favour of the reduced model (null hypothesis; PCI). Additionally, Supplementary Figure S5 displays a matrix of scatterplots comparing questionnaire scores and task variables from the RMCT across the discovery and the validation data set (*N*_tot_=50).

## Discussion

In this work, we propose a concrete process model for perceived control over breathing perturbations. After developing a novel behavioural paradigm, the RMCT, that allows for perturbations of perceived control over inspiratory resistive loads, we used data from healthy volunteers to test our proposed model. We found ‘very strong’ evidence in favour of our primary hypothesis that perceived control over breathing perturbations can be explained as a function of interoceptive prediction errors in the RMCT. The model that provided the best explanation of the perceived control over breathing perturbation rating data in the RMCT was an autoregressive model of order 1 which used a combination of interosensory inputs *u* (absence/presence of inspiratory resistive loads) and signed prediction errors *δ* which encode how unexpected these inputs were. In our model, we found a positive effect of interosensory inputs *u* (normal breathing=1 and restricted breathing=-1) and a negative effect of signed PEs *δ* on ratings of perceived control over breathing perturbations. Regarding our additional hypotheses, we found ‘positive’ evidence for an association between the weighting parameter *γ* of past experience of control over breathing perturbations and fatigue (MFIS). All other hypothesis tests regarding associations between task-based variables and psychological questionnaire scores remained inconclusive.

Concerning our primary hypothesis, Bayesian model selection revealed that the full model ***M****_δ_*_,*u*_, incorporating both interosensory inputs *u* and signed PEs *δ* alongside an autoregressive component, outperformed a null model and two reduced autoregressive model variants. Regarding our additional hypotheses, we found ‘positive’ evidence for an association between MAP estimates of the computational model parameter *γ* and fatigue (MFIS). In other words, the more strongly an individual’s current estimate of perceived control over breathing perturbations depended on past experiences of control, the higher the associated level of fatigue,. All other additional hypothesis tests investigating associations between task-based variables and other psychological questionnaire scores yielded ‘barely worth mentioning’ levels of evidence either for or against the null hypothesis (Kass and Raftery, 1995). This indicates that while some links between these different variables and constructs might exist, our current data set does not provide sufficient evidence for robust conclusions. Our sample size of *N*_vs_=30 was determined by our primary hypothesis, using a Sequential Bayes Factor design (Schönbrodt et al., 2017), not by the additional hypotheses 2-5. This may have led to a sample size with insufficient power for detecting links between model parameters and questionnaire scores.

The question of this study was motivated by the fact that the perception of control is an important determinant of mental health (see Introduction). While different forms of perceived control exist, perceived control over the regulation of bodily states has taken a particularly important role in computational frameworks of mental health (Petzschner et al., 2017), including the allostatic self-efficacy (ASE) theory of fatigue (Stephan et al., 2016). Experimental investigations of perceived control over bodily states have mainly concerned the domain of pain (e.g. Habermann and Büchel, 2025; Wiech et al., 2006). By contrast, investigating perceived control over other bodily states experimentally is more challenging, and we are not aware of previous work that concerns processes in classical interoceptive domains such as perception of cardiac, respiratory, or intestinal processes. Notably, as implied by contemporary predictive processing concepts of brain-body interactions (Gu et al., 2013; Petzschner et al., 2017; Pezzulo et al., 2015; Seth and Friston, 2016; Smith et al., 2017; Stephan et al., 2016), control over the regulation of bodily states can be instantiated in multiple ways, including: (i) homeostatic/allostatic control in terms of explicit behavioural actions that affect bodily states, e.g. by altering the body’s exposure to environmental perturbations; (ii) homeostatic control via reflex arcs that act on ascending viscerosensory signals and elicit descending neuronal and/or hormonal actions; (iii) updating of beliefs that underlie allostatic forecasting and metacognitive evaluations, respectively. The experimental paradigm in this study used a strategy that targeted specifically the first of these different forms of exerting control: by learning the visuomotor skills required by our gamified task, participants gained a degree of control over whether they were exposed to aversive interosensory inputs, i.e. whether or not breathing was impeded by resistive loads.

Our primary hypothesis test could be mistaken for a direct test of the ASE theory. This is not the case, for at least two reasons. First, the restriction of our paradigm to explicit behavioural control over the exposure to environmental perturbations (see previous paragraph) means that our study covers only a subpart of the homeostatic/allostatic control processes considered by the ASE theory. Second, our paradigm uses only short-lived perturbations of homeostasis and manipulates the perception of control on a short time scale of seconds to minutes; by contrast, the ASE theory refers to a chronic loss of perceived control over bodily states, induced by a lasting state of systemic dyshomeostasis which is reflected by continuously elevated interoceptive PE levels and is immune to behavioural and autonomic attempts of restoring homeostasis.

Having said this, our model of perceived control is motivated by and has strong conceptual links to the ASE theory. For example, as in the ASE theory, our model describes a metacognitive process operating on a cognitive (algorithmic) variable, i.e. viscerosensory PE, and explains fluctuations in the participants’ experience of allostatic self-efficacy or perceived control over bodily states. This also provides a natural link between our work and research on metacognition and self-efficacy. Research on metacognition has mostly focused on self-evaluation of memory, perception, and decision-making (Fleming and Dolan, 2012). Our work represents, to the best of our knowledge, the first empirical investigation of metacognition in the domain of allostatic control. Hence, it is not straightforward to compare our model to existing models of metacognitive belief formation. However, an important characteristic inherent to different models of metacognition which our current process model of allostatic self-efficacy belief formation is missing is an explicit representation of (first-order) actions (Fleming and Daw, 2017). In other words, our model of perceived control over breathing perturbation is currently agnostic of our participants’ task behaviour, i.e. the selected jump strength and the avatar’s landing position (including its distance from the target island). Fleming and Daw (2017) proposed “a general Bayesian framework in which self-evaluation is cast as a ‘second-order’ inference”, which postulates that one’s actions carry important information contributing to self-evaluation. This second-order inference framework predicts that prospective metacognitive judgements should be less accurate than retrospective judgements, which is in line with empirical findings (Siedlecka et al., 2016). Based on these considerations, a natural extension of our model of allostatic self-efficacy belief formation would be an explicit representation of task behaviour and its role in self-efficacy belief formation.

It is also worth mentioning that the reparameterised form of our model (Eq. 2) shares a degree of structural resemblance with a computational model of an unrelated construct, i.e. momentary subjective well-being (Rutledge et al., 2014). The similarity is that in the model by Rutledge et al., PEs – albeit of a different kind – are also thought to contribute to temporal fluctuations of subjective experience. In brief, their model explained happiness ratings as a combination of certain rewards, the expected values of chosen gambles, and reward PEs. However, otherwise the two models have little in common as they are conceptually rooted in different concepts (predictive processing vs. reinforcement learning theories).

Our study has several additional limitations. First, while sample size was determined by our primary hypothesis, it limited our ability to draw firm conclusions about our secondary hypotheses. Second, we specifically investigated perceived control in the domain of breathing. It is unclear whether our findings generalise to other domains of interoception. Third, the construct of control as operationalised by our task, does not fully capture the multifaceted nature of control over bodily states, as discussed above. Finally, while carefully designed, the ecological validity of the task is inherently constrained by its laboratory setting, potentially limiting the generalisability of our findings to real-world scenarios of perceived control over breathing or bodily states in general.

Having said this, our study also possesses several notable strengths. First, we developed a process model of perceived control over breathing perturbations which, to our knowledge, is the first generative model which describes a dynamic metacognitive processes of how the brain evaluates its capacity for control over bodily states and can be applied to simple behavioural readouts. The model parameters have clear interpretations and are well identifiable, as shown by parameter recovery analyses. Second, model development harnessed a principled Bayesian workflow (Hess et al., 2025) and used separate discovery and validation datasets. Third, sample size was determined in a principled manner, using a Sequential Bayes Factor design. Fourth, in order to maximise robustness, transparency and reproducibility, the entire analysis pipeline was pre-registered, with all deviations from the analysis plan clearly marked in this manuscript, and underwent independent code review (Pereira et al., 2026).

In summary, we have presented a novel, validated process model of perceived control over breathing perturbations which can be applied to simple behavioural measures. This model may find useful application in computational psychiatry investigations, particularly of disorders where the dynamics of perceived loss of control over bodily states corresponds to the timescale of perturbations in the RMCT task. We hope that this model will provide a starting point for further development of models assessing metacognition of homeostatic/allostatic control and will support the development of computational assays for clinical applications (Stephan and Mathys, 2014).

## Authors’ contributions

AJH, conceptualisation, data curation, formal analysis, investigation, project administration, methodology, software, visualisation, writing – original draft, writing – review and editing, publication of data and code; GC, LD, conceptualisation, methodology, software, writing – review and editing; SMe, code review, writing – review and editing; SI, conceptualisation, data curation, project administration, methodology, supervision, writing – review and editing; LK, SMa, CB, AK, data curation, investigation, writing – review and editing; CM, SF, JS, OKH, JH, conceptualisation, methodology, writing – review and editing; KES, resources, conceptualisation, methodology, funding acquisition, supervision, writing – review and editing.

## Acknowledgements

We would like to thank Matthias-Müller Schrader, Dina von Werder, Lilian Weber, Ryan Smith and Katherina Hemmo for helpful discussions and feedback, Imre Kertesz and Sebastian Grässli for help with experimental setup in the laboratory, and Natalie Araya for support regarding ethics approval application and administration. We thank Noé Zimmermann, Jana Bünzli, Patrizia Sting, and Nathalie Appenzeller for their help with recruitment, data checks, and research database administration.

