## Supplementary Materials for "Perceived Control Over Breathing Perturbations Depends on Interosensory Input and Prediction Error"

to

### S1. Parameter Recovery

Figures S1A-D show results from parameter recovery analyses of all four models.

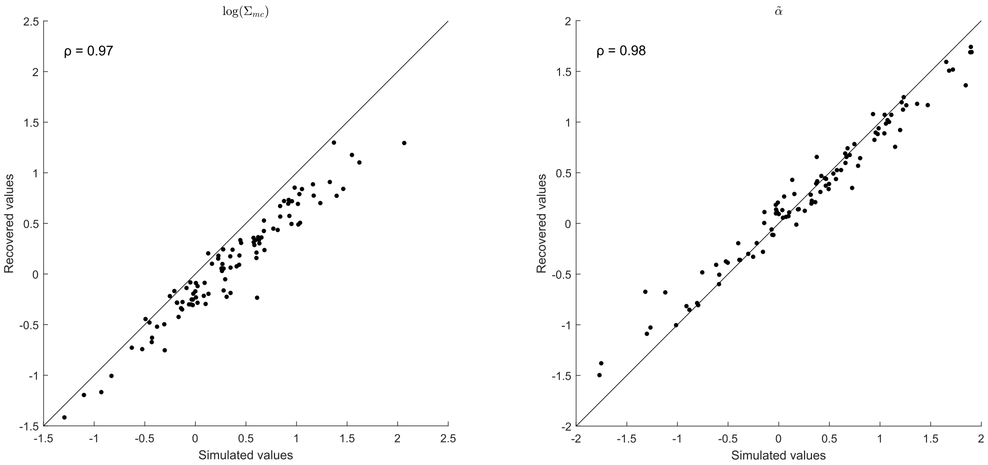


Figure S1A | Parameter recovery of $\boldsymbol{M_0}$. Displayed are simulated against estimated parameter values ($N_{sim} = 100$) as well as Pearson correlation coefficients ($\rho$). The identity line (black) represents perfect recovery.

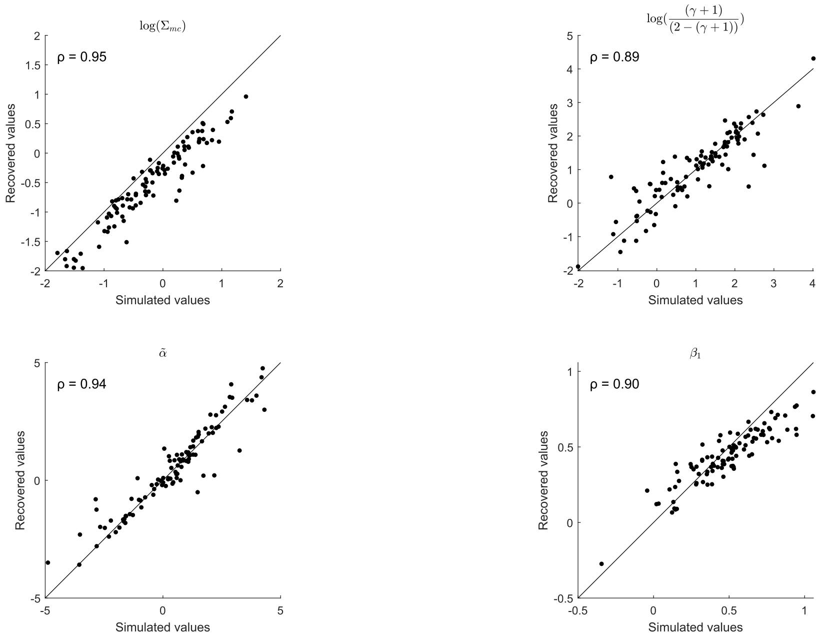


Figure S1B | Parameter recovery of $\boldsymbol{M_u}$. Displayed are simulated against estimated parameter values ($N_{sim} = 100$) as well as Pearson correlation coefficients ($\rho$). The identity line (black) represents perfect recovery.

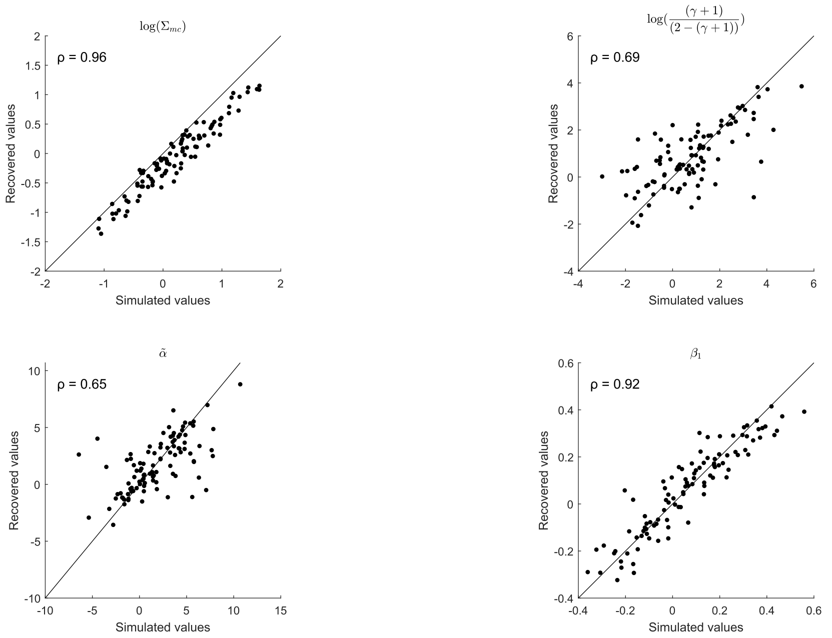


Figure S1C | Parameter recovery of $\boldsymbol{M_{\delta}}$. Displayed are simulated against estimated parameter values ($N_{sim} = 100$) as well as Pearson correlation coefficients ($\rho$). The identity line (black) represents perfect recovery.

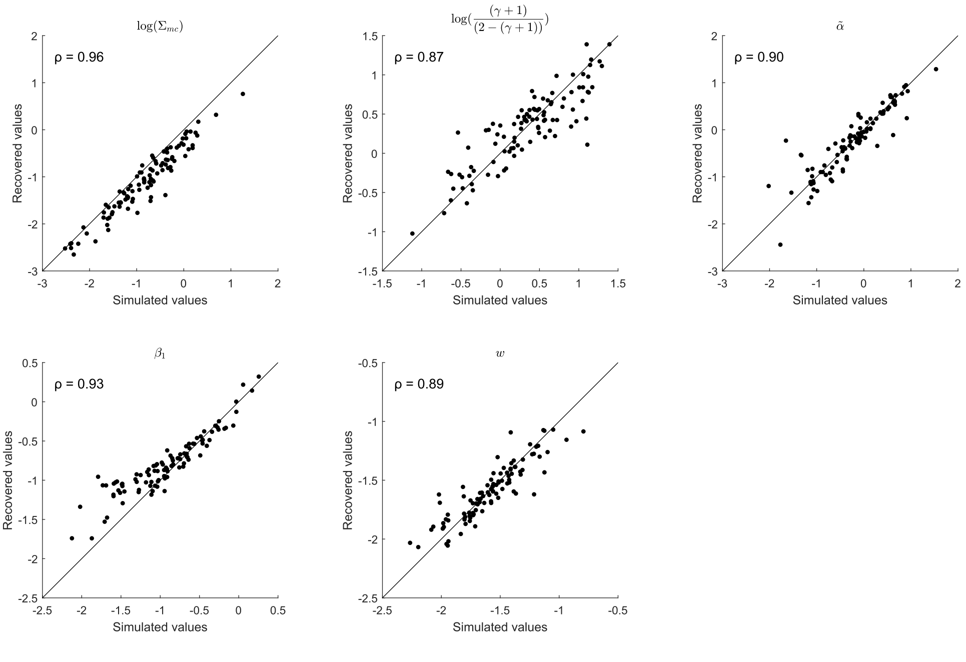


Figure S1D | Parameter recovery of $\boldsymbol{M_{\delta,u}}$. Displayed are simulated against estimated parameter values ($N_{sim} = 100$) as well as Pearson correlation coefficients ($\rho$). The identity line (black) represents perfect recovery.

S2. *Model Identifiability*

Figure S2A shows confusion matrices resulting from model identifiability analyses.

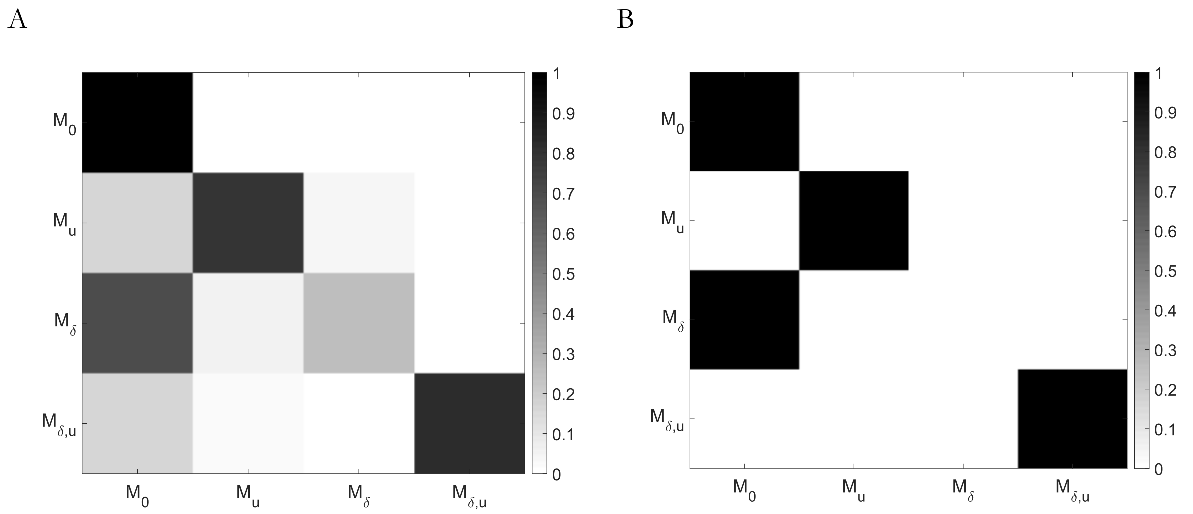


Figure S2A | Model identifiability confusion matrices. Simulated models are displayed on the x axis, recovered models on the y axis. **A** shows frequencies of highest approximate log model evidence (LME) values, **B** shows protected exceedance probabilities (PXP) from random-effects Bayesian model selection (RFX BMS).

Our models showed excellent identifiability apart from $\boldsymbol{M_\delta}$. Apparent non-identifiability of $\boldsymbol{M_\delta}$ was a consequence of the Bayesian nature of our simulation analyses and the use of the discovery set to elicit *empirical priors* in a regime of actual human behaviour. As shown in Supplementary Figure S3C, the *empirical prior* density of the $\beta_1$ parameter in model $\boldsymbol{M_\delta}$ is tightly concentrated around 0 (mean 0.05, variance 0.05). As a consequence, most of the sampled $\beta_1$ values are close to 0 and the simulated behaviour thus resembles that of our null model ($\boldsymbol{M_0}$). Supplementary Figure S2B shows that for synthetic responses generated by $\boldsymbol{M_\delta}$ using $\beta_1$ values close to 0, $\boldsymbol{M_0}$ provides the best explanation for the data (highest approximate LME values), whereas for synthetic responses generated by $\boldsymbol{M_\delta}$ using $\beta_1$ values further away from 0, $\boldsymbol{M_\delta}$ explains the data best in most cases. Hence, there is no reason to doubt the identifiability of models. Assessing the identifiability of the full model ($\boldsymbol{M_{\delta,u}}$) and the null model ($\boldsymbol{M_0}$) based on the log BF distribution on the respective synthetic data sets confirms this intuition (Supplementary Figure S2C).

Figure S2B displays an in-depth analysis of the identifiability of $\boldsymbol{M_\delta}$ by displaying models with highest approximate log model evidence (LME) values against $\beta_1$ parameter values of simulated subjects generated using $\boldsymbol{M_\delta}$.

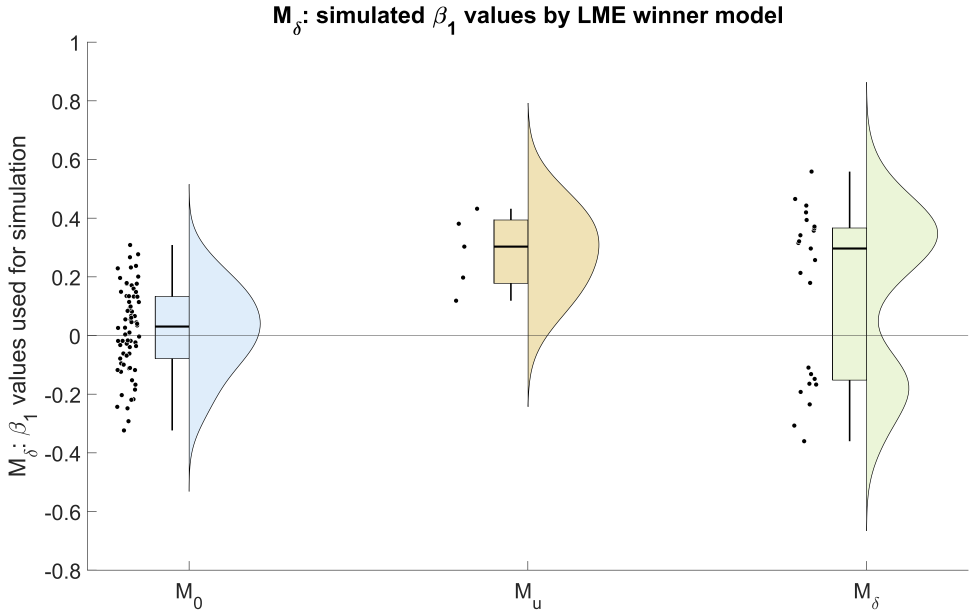


Figure S2B | Detailed model identifiability analysis of $\boldsymbol{M}_{\delta}$. Displayed are models with highest approximate log model evidence (LME) values against $\boldsymbol{\beta}_1$ parameter values used for generating data under $\boldsymbol{M}_{\delta}$.

Figure S2C shows log Bayes Factor (BF) distributions between the full model ($\boldsymbol{M}_{\delta,u}$) and the null model ($\boldsymbol{M}_0$) on the respective synthetic data sets.

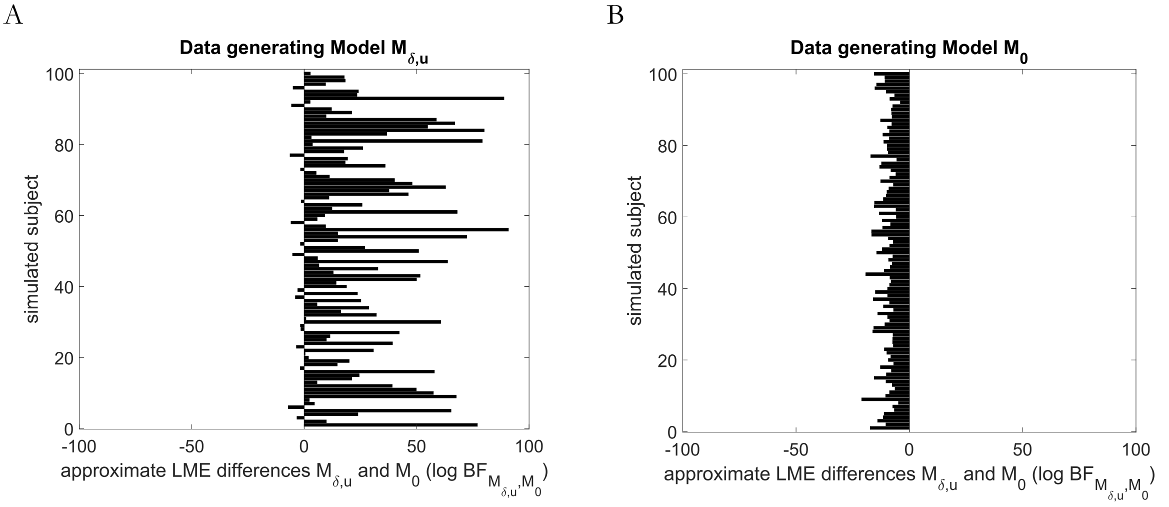


Figure S2C | Log Bayes Factor (BF) distribution between the full model ($\boldsymbol{M}_{\delta,u}$) and the null model ($\boldsymbol{M}_0$) on the respective synthetic data sets. **A** shows log BFs calculated on the synthetic data set generated by $\boldsymbol{M}_{\delta,u}$ and **B** for data generated by $\boldsymbol{M}_0$, respectively.

#### S3: Priors

Figures S3A-D show *initial* and *empirical prior* densities as well as MAP estimates on the discovery set ($N_{ds} = 20$) for all four models.

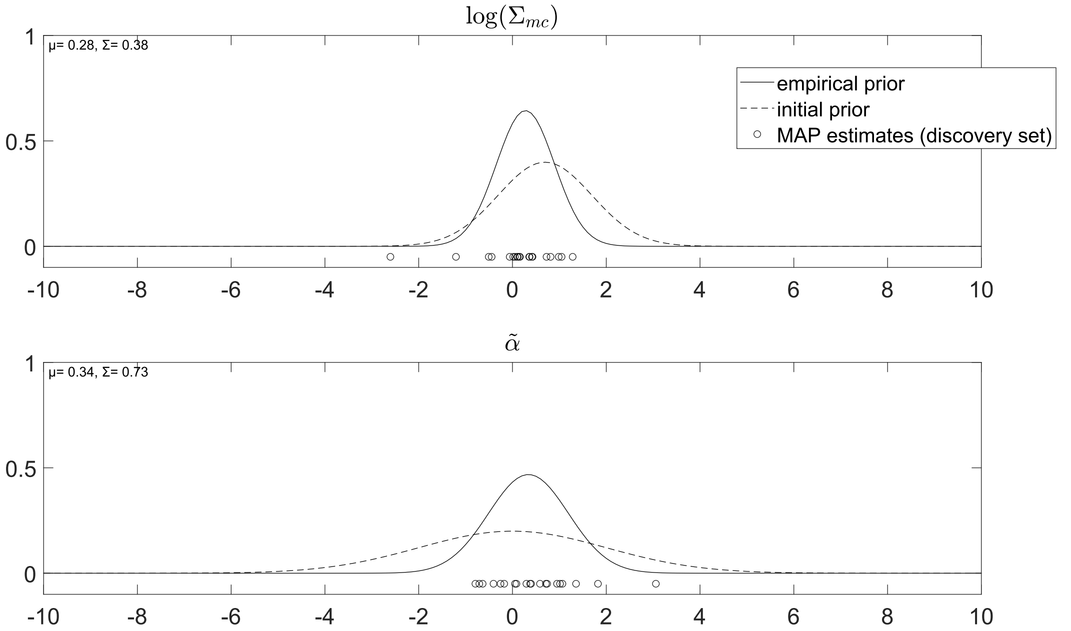


Figure S3A | *Initial* and *empirical* prior densities as well as MAP estimates for all free parameter of $\boldsymbol{M}_0$ resulting from model inversion on the discovery set ($N_{ds} = 20$). Sufficient statistics ($\mu, \Sigma$) of the estimated *empirical* prior density for each parameter are indicated in the top left corner of each subplot.

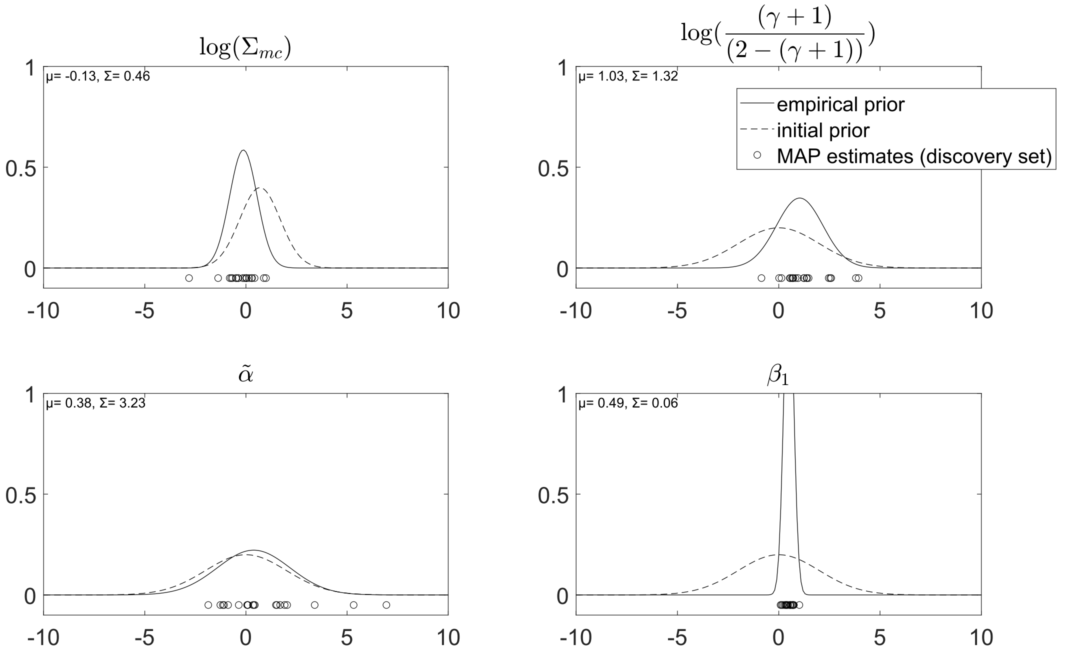

Figure S3B | *Initial* and *empirical* prior densities as well as MAP estimates for all free parameter of $\boldsymbol{M}_u$ resulting from model inversion on the discovery set ($N_{ds} = 20$). Sufficient statistics ($\mu, \Sigma$) of the estimated *empirical* prior density for each parameter are indicated in the top left corner of each subplot.

63

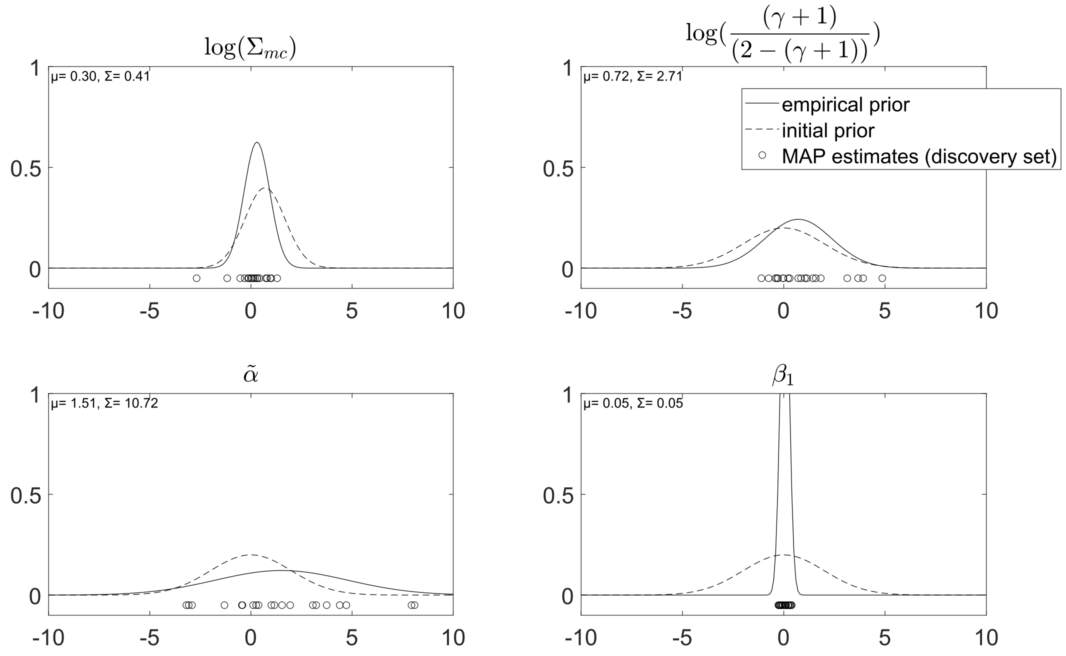


Figure S3C | *Initial* and *empirical* prior densities as well as MAP estimates for all free parameter of $\boldsymbol{M}_\delta$ resulting from model inversion on the discovery set ($N_{ds} = 20$). Sufficient statistics ($\mu, \Sigma$) of the estimated *empirical* prior density for each parameter are indicated in the top left corner of each subplot.

64

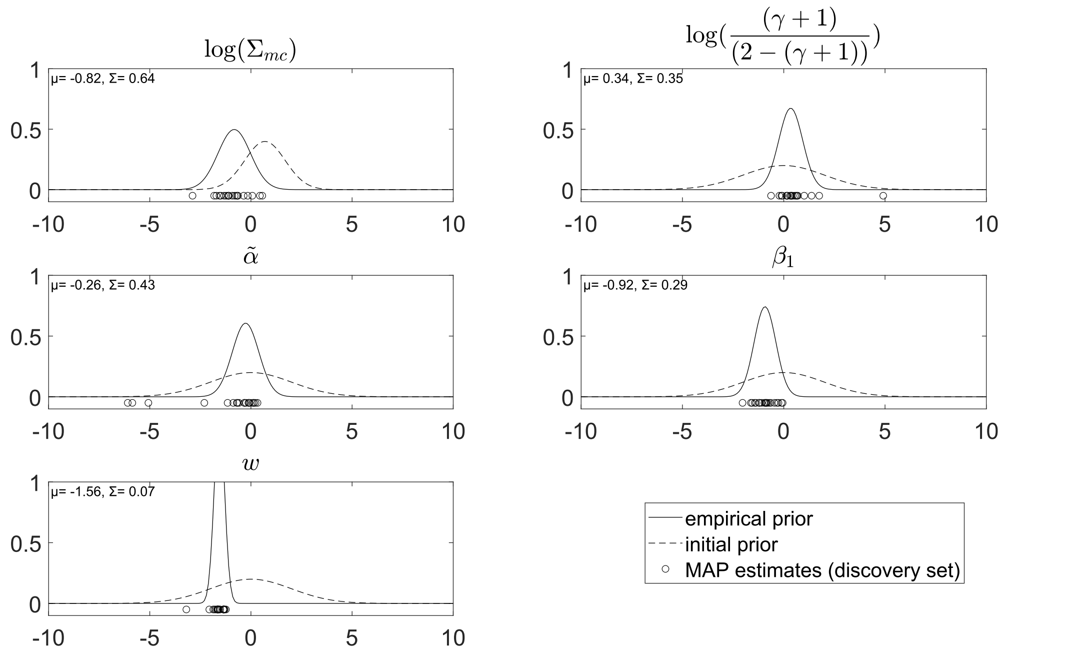


Figure S3D | *Initial* and *empirical* prior densities as well as MAP estimates for all free parameter of $\boldsymbol{M}_{\delta,u}$ resulting from model inversion on the discovery set ($N_{ds} = 20$). Sufficient statistics ($\mu, \Sigma$) of the estimated *empirical* prior density for each parameter are indicated in the top left corner of each subplot.

65

#### S4: Model Evaluation

Figures S4A-B show example single subject fits across all four models of the best and worst fitting subject, respectively, evaluated based on explained variation (coefficient of determination of $\boldsymbol{M}_{\delta,u}$). Quantitatively, model fit for individual ranged from $R^2 = 0.15$ (worst model fit) to $R^2 = 0.93$ (best model fit) across our sample.

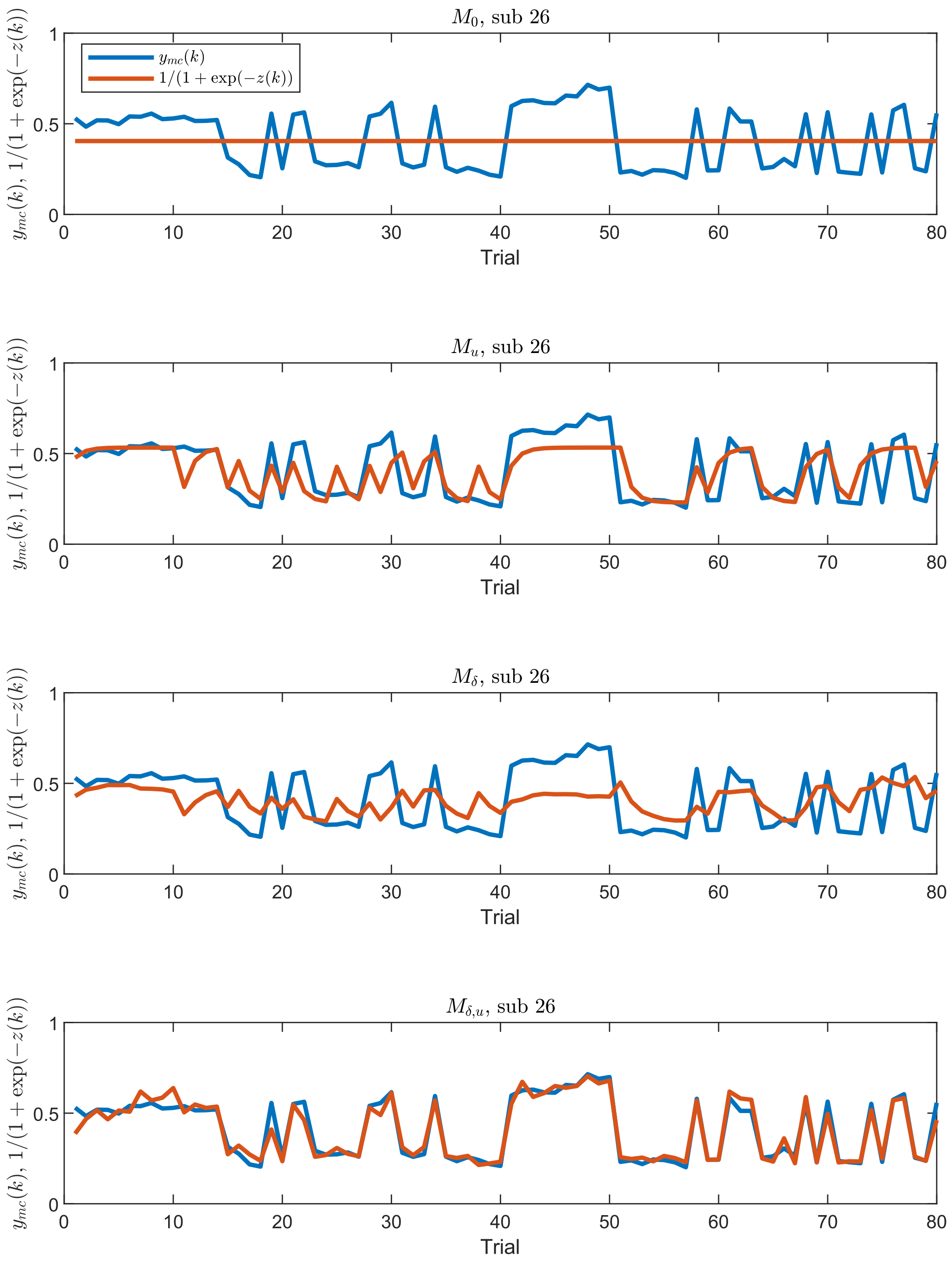


Figure S4A | Individual model fits for best fitting participant in the validation set ranked according to the coefficient of determination of $\boldsymbol{M}_{\delta,u}$.

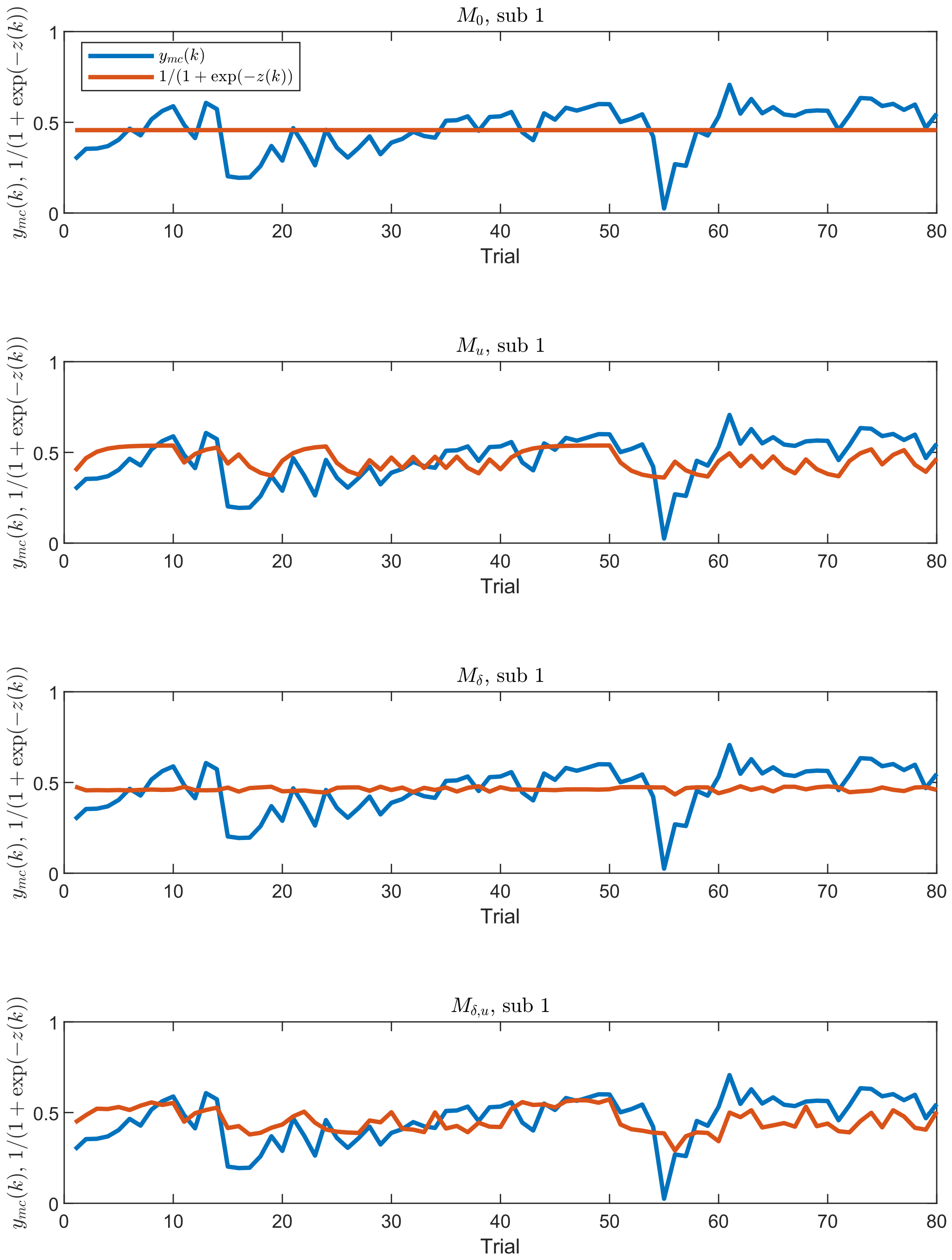


Figure S4B | Individual model fits for worst fitting participant in the validation set ranked according to the coefficient of determination of $\boldsymbol{M}_{\delta,u}$.

Figure S4C displays an example single participant fit from the validation set across all four models (upper panel A) alongside a didactic overview of the decomposed predicted perceived control over breathing perturbation ratings for the recursive formulation of $\boldsymbol{M}_{\delta,u}$ (lower panel B).

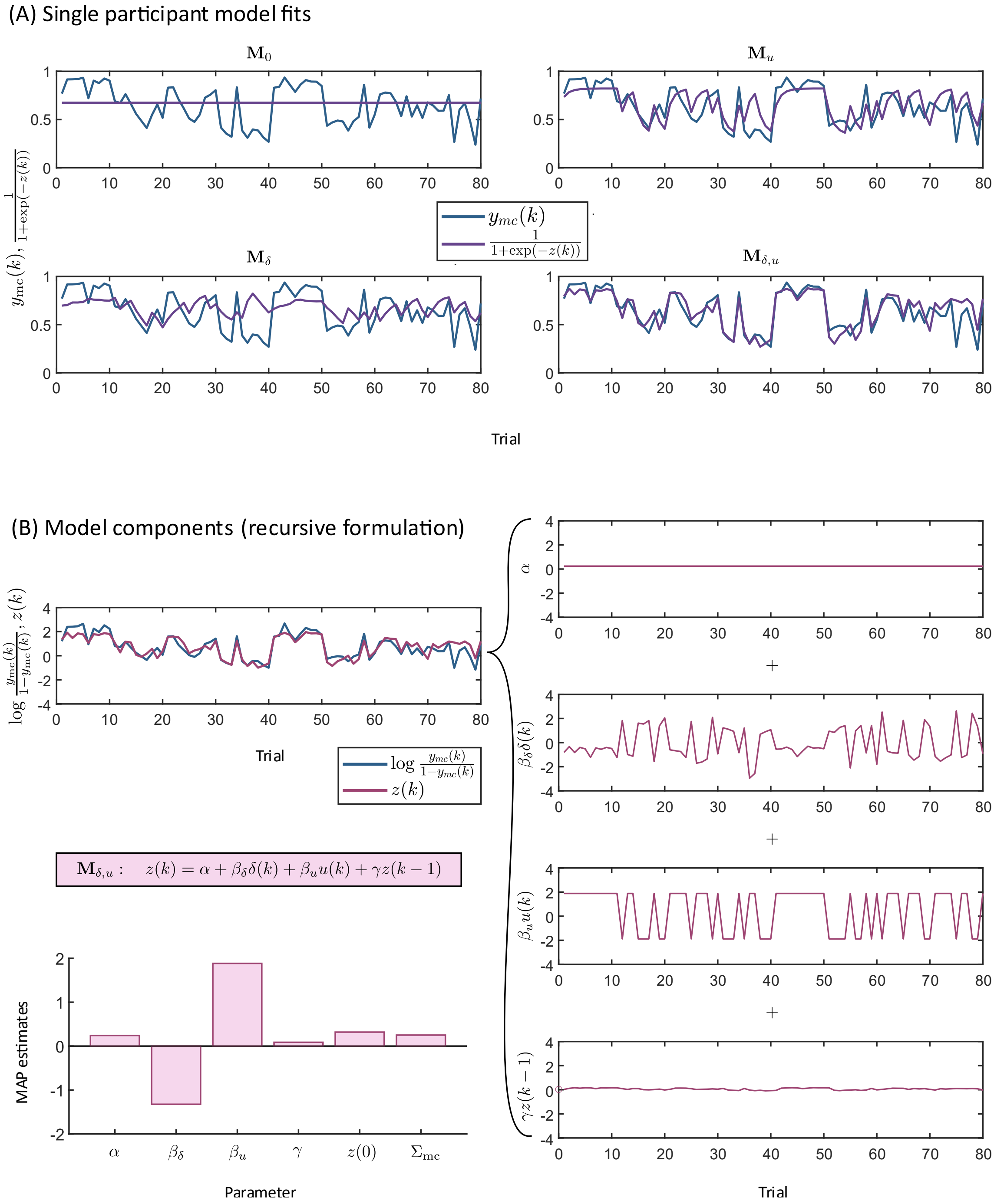


Figure 5 | Single participant data from the validation set and model fits. (A) Perceived control over breathing perturbation ratings $y_{\text{mc}}(k)$ of a single participant (blue) and predicted ratings $\frac{1}{1+\exp(-z(k))}$ for all four models (purple). (B) Perceived control over breathing perturbation ratings $\log \frac{y_{\text{mc}}(k)}{1-y_{\text{mc}}(k)}$ of a single participant in estimation space (blue) and predicted ratings $z(k)$ for the full model $\boldsymbol{M}_{\delta,u}$, including decomposed components of the AR(1) model (recursive formulation, Eq. 1 in the main manuscript) in the right panel. In the lower left corner, MAP estimates of all parameters for this participant are visualised in a barplot.

#### S5 Additional Hypotheses

Our additional hypotheses concerned the question how task-based readouts from the RMCT are related to constructs of fatigue and control that were measured by established questionnaires. In our statistical analysis plan, we specified that we would perform Bayesian ANCOVAs for different dependent variables (questionnaire scores) on the validation set ($N_{vs} = 30$). For each ANCOVA model in H2-H5, we included only a subset of the previously described predictors. The choice of subset was motivated by exploratory analyses on the discovery set ($N_{ds} = 20$), which was also pre-specified in our statistical analysis plan. Specifically, we selected those predictors for whom the odds for inclusion was greater after ($\frac{P(incl|D)}{1-P(incl|D)}$) than before seeing the data ($\frac{P(incl)}{1-P(incl)}$) (Bergh et al., 2020).

For example, Hypothesis 2 (H2) proposed that a subset from all model-based estimates of perceived control over breathing perturbation ($\gamma, w, \beta_1, \tilde{\alpha}$), other ASE-inspired readouts (mean and variances of overall control and tolerance ratings; $\bar{\mathbf{y}}_c$, $Var(\mathbf{y}_c)$, $\bar{\mathbf{y}}_{tol}$, $Var(\mathbf{y}_{tol})$), and task unpleasantness ($y_{av}$) in the RMCT explained overall fatigue scores from standard questionnaires (FAS, MFIS). The MFIS score was therefore selected as the dependent variable in the Bayesian ANCOVA. As statistical test, we prespecified the Bayes factor $BF_{10}$, comparing the full model $M_{1,H2}$ containing the entire subset of predictors ($\gamma$, $\bar{\mathbf{y}}_c$, $Var(\mathbf{y}_{tol})$, $y_{av}$, gender, age, intercept) against reduced model $M_{0,H2}$ containing only gender, age, and an intercept. As exploratory analyses, we computed Bayes factors for all nested models of $M_{1,H2}$ against $M_{0,H2}$. Additionally, we conducted a sensitivity analysis using the FAS instead of the MFIS score as dependent variable together with the subset of predictors identified on the discovery set.

As described in the main text, the process of internal code review upon completion of the analyses revealed an error in the scoring of the PSQI questionnaire as well as an indexing error in the analysis of the questionnaire data leading to participants being associated with incorrect data. As a consequence, we recomputed the entire analysis including the selection of independent variables on the discovery set, which resulted in a different subset of predictors for H2-H5 compared to those listed in the analysis plan. The results from the corrected version of the analysis and the updated set of predictors for the different Bayesian ANCOVAs are reported in Supplementary Table 1. For those dependent variables where the re-analysis yielded no relevant predictors on the discovery set, we did not conduct any tests on the validation set and the respective rows in Supplementary Table 1 are left empty.

*Supplementary Table 1* | Results of the analyses for hypotheses 2-5. Displayed are the dependant variables (Dep. Var.) for each Bayesian ANCOVA of hypotheses 2-5 (Hyp $i$). $BF_{10}$ denotes the Bayes Factor comparing $M_{1,Hi}$ (covariates additional to $M_{0,Hi}$ listed in the respective column) and $M_{0,Hi}$ (covariates and factors additional to an intercept term listed in the respective column). $BF_{best,0}$ denotes the Bayes Factor comparing the best nested model of $M_{1,Hi}$ ($M_{best,Hi}$; covariates additional to $M_{0,Hi}$ listed in the respective column) and $M_{0,Hi}$. Bayes Factors are colour coded according to the classification by Kass and Raftery (1995) with evidence in favour of the alternative hypothesis in shades of orange and evidence in favour of the null hypothesis in shades of blue. Empty rows represent dependent variables where we did not conduct any hypothesis tests on the validation set because the required re-analysis due to the discovered coding error yielded no relevant predictors on the discovery set.

| Hyp $i$ | Dep. Var. | $M_{1,Hi}$ | $M_{0,Hi}$ | $BF_{10}$ | $M_{\text{best},Hi}$ | $BF_{best,0}$ |
|---|---|---|---|---|---|---|
| $i = 2$ | MFIS | | | | | |
| | FAS | $y_{av}$ | age, gender | 1.617 | $y_{av}$ | 1.617 |
| $i = 3$ | MFIS_cog | | | | | |
| | MFIS_phys | | | | | |
| | MFIS_psychosoc | $Var(\mathbf{y}_{tol}), y_{av}$ | age, gender | 0.325 | $y_{av}$ | 0.599 |
| $i = 4$ | MFIS | $\gamma, Var(\mathbf{y}_{tol})$ | MAIA-2_38, PSQI, age, gender | 2.573 | $\gamma$ | 3.231 |
| | FAS | | | | | |
| $i = 5$ | GSES | $y_{av}$ | age, gender | **37.572** | $y_{av}$ | **37.572** |
| | PCI | $\bar{\mathbf{y}}_c, \tilde{\alpha}$ | age, gender | 0.280 | $\bar{\mathbf{y}}_c$ | 0.502 |
| | LOC_p | $\bar{\mathbf{y}}_c, Var(\mathbf{y}_c)$ | age, gender | 1.379 | $\bar{\mathbf{y}}_c, Var(\mathbf{y}_c)$ | 1.379 |
| | LOC_c | $\bar{\mathbf{y}}_c, Var(\mathbf{y}_c)$ | age, gender | 1.838 | $\bar{\mathbf{y}}_c, Var(\mathbf{y}_c)$ | 1.838 |

During internal discussions of the analyses for our additional hypotheses, we reached a consensus that Bayesian Linear Regression (BLR) would be more appropriate as statistical tool to test our hypotheses compared to Bayesian ANCOVA. ANCOVA examines differences in group means while controlling for covariates, whereas Linear Regression predicts a continuous outcome from predictors including covariates. Our linear models consisted of continuous variables only (except the covariate gender). Additionally, our goal was to understand how predictors related to the dependent variable and not a comparison between groups after controlling for covariates.

Therefore, we decided to re-compute the entire analysis for our additional hypotheses, including the selection of predictors on the discovery set following the same procedure as described above. The results of the Bayesian Linear Regression analyses are reported in Table 3 of the main manuscript. Notably, the selection of predictors on the discovery set yielded fewer predictors for whom the $BF_{incl} > 1$, i.e. the odds for inclusion was greater after than before seeing the data compared to the Bayesian ANCOVA setting. This seems odd at first glance, can however be explained by the different type of priors implemented in JASP for Bayesian ANCOVA as opposed to BLR. In other words, both procedures yield the same order of predictors when ranked according to the $BF_{incl}$, but the value of the $BF_{incl}$ changes depending on the choice of prior.

In Supplementary Table 2, we report the full results of the BLR model comparison for the dependent variable MFIS (H4) in the form of the JASP output. The posterior summary of coefficients resulting from the same analysis are listed in Supplementary Table 3 in the form output by JASP.

*Supplementary Table 2* | Model comparison results of the Bayesian Linear Regression using MFIS as the dependent variable and $\gamma$, age, gender, MAIA-2_38, and PSQI as dependent variables.

| Models | P(M) | P(M\|data) | $BF_M$ | $BF_{10}$ | $R^2$ |
|---|---|---|---|---|---|
| Null model (incl. age, gender, MAIA38, PSQI) | 0.500 | 0.243 | 0.320 | 1.000 | 0.146 |
| gamma | 0.500 | 0.757 | 3.122 | 3.122 | 0.300 |

*Note.* All models include age, gender, MAIA38, PSQI.

*Supplementary Table 3* | Posterior Summaries of Coefficients of the Bayesian Linear Regression using MFIS as the dependent variable and $\gamma$, age, gender, MAIA-2_38, and PSQI as dependent variables.

| | | | | | | | | 95% Credible Interval | |
|---|---|---|---|---|---|---|---|---|---|
| Coefficient | P(incl) | P(excl) | P(incl\|data) | P(excl\|data) | $BF_{inclusion}$ | Mean | SD | Lower | Upper |
| Intercept | 1.000 | 0.000 | 1.000 | 0.000 | 1.000 | 25.000 | 2.587 | 19.592 | 30.486 |
| age | 1.000 | 0.000 | 1.000 | 0.000 | 1.000 | 0.167 | 0.534 | -0.985 | 1.239 |
| gender | 1.000 | 0.000 | 1.000 | 0.000 | 1.000 | 5.869 | 4.380 | -3.271 | 15.003 |
| gamma | 0.500 | 0.500 | 0.757 | 0.243 | 3.122 | 14.609 | 12.168 | -1.552 | 38.999 |
| MAIA38 | 1.000 | 0.000 | 1.000 | 0.000 | 1.000 | -1.438 | 1.596 | -4.785 | 1.857 |
| PSQI | 1.000 | 0.000 | 1.000 | 0.000 | 1.000 | 0.733 | 1.283 | -1.935 | 3.409 |

Figure S5 displays a matrix of scatterplots comparing questionnaire scores and task variables from the RMCT across the discovery and the validation data set ($N_{\text{tot}} = 50$).

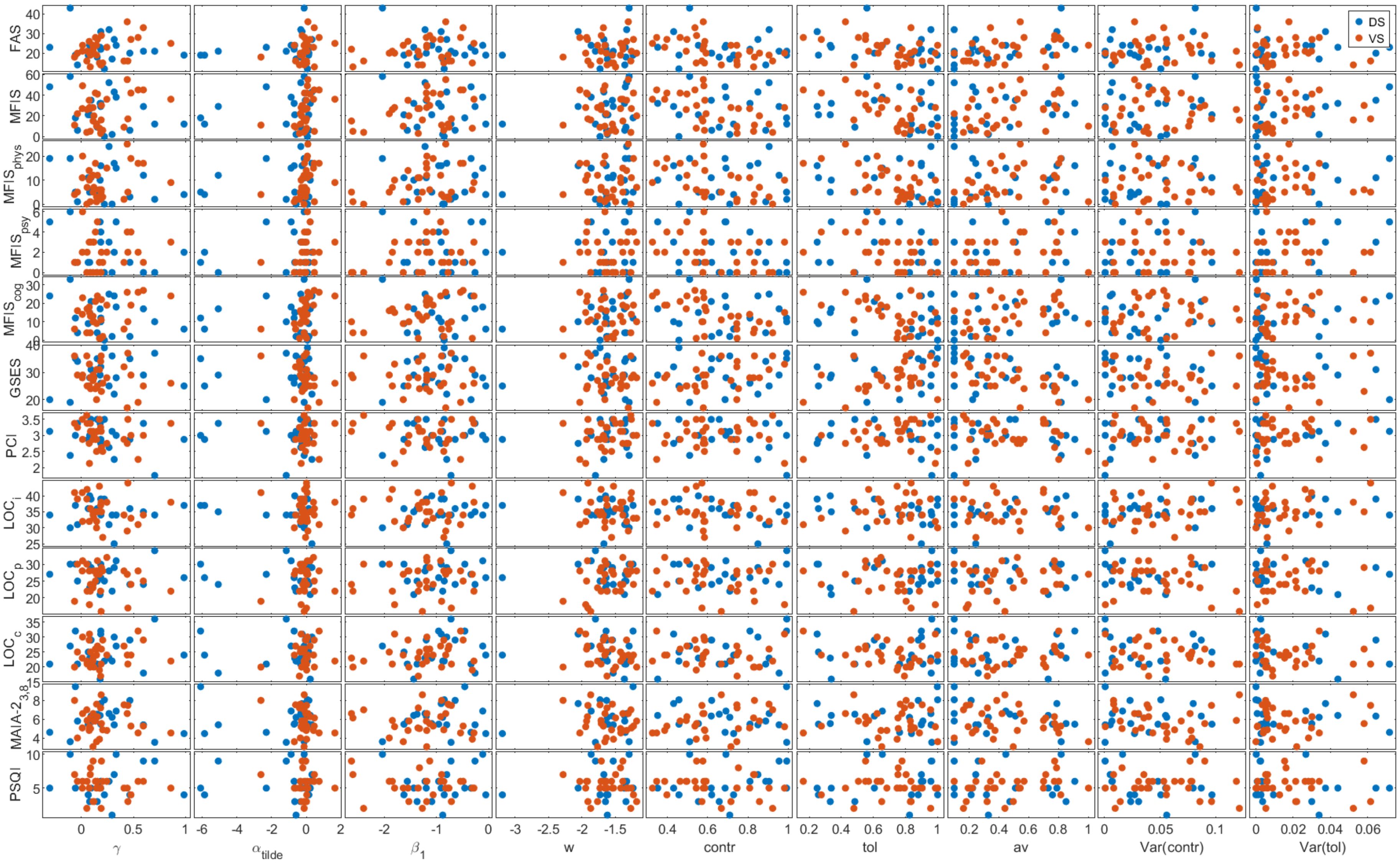


Figure S5 | Matrix of scatterplots comparing questionnaire scores (y axis) and task variables from the RMCT (x axis) across all participants ($N_{tot} = 50$) colour-coded by discovery (blue) and validation (red) data set.

#### S6 Breathing Circuit

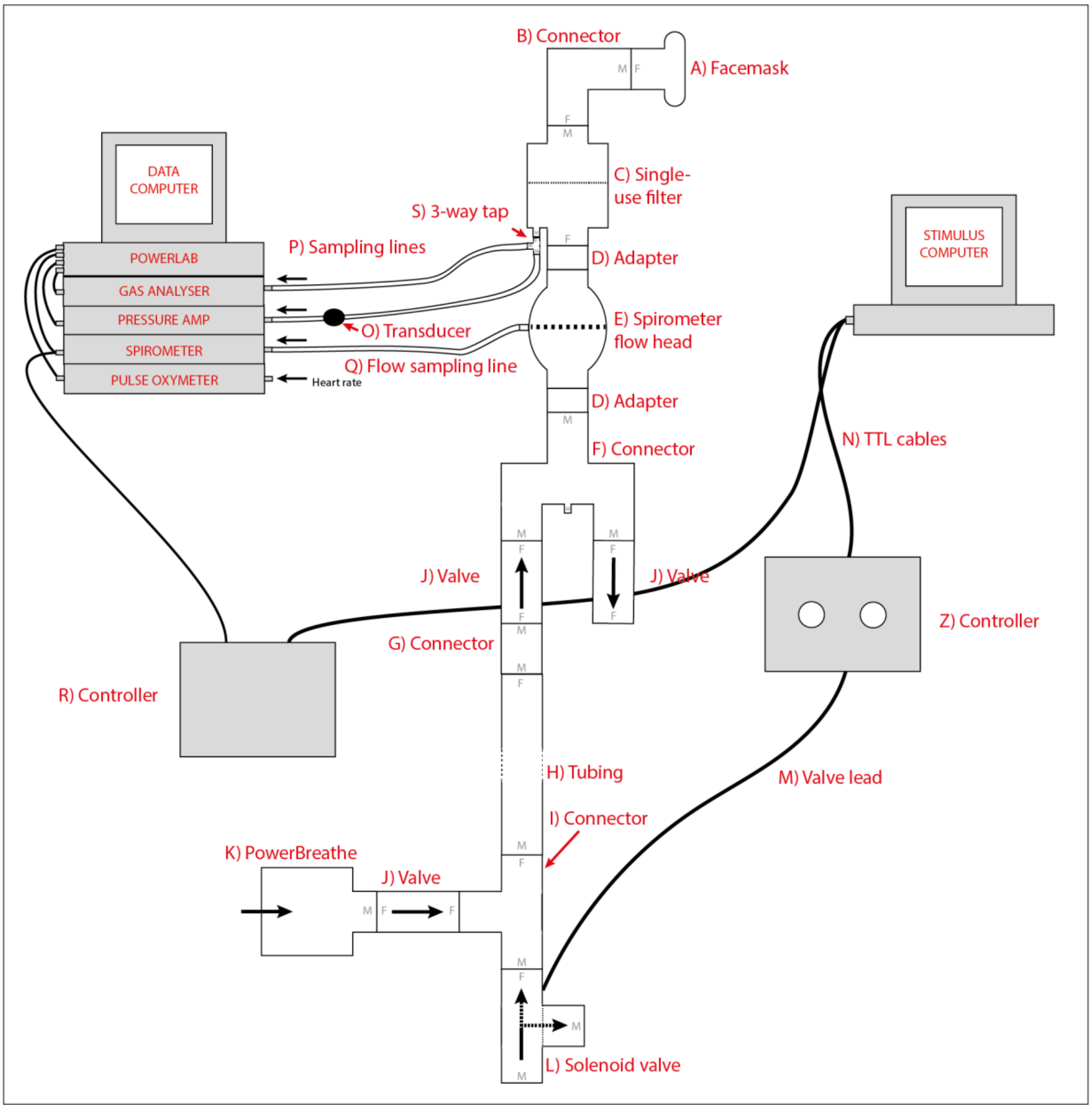


Figure S6 | A schematic overview over the breathing circuit used for administration of inspiratory resistive loads and physiological measurements. Participants were connected to the mechanical breathing system via a facemask covering their mouth and nose which was attached to a bacterial and viral filter (labelled C). Sampling lines to a pressure transducer (labelled O) and amplifier was used to record inspiratory pressure and to a gas analyser for respiratory gases. A spirometer connected to a respiratory flow head (labelled E) was used to measure inspiratory flow. Inspiratory resistive loads were induced automatically triggered by the stimulus computer via controller box (labelled Z) and solenoid valve (labelled L) to redirect the supply of unobstructed air from the environment to the PowerBreathe device (labelled K), which provided a constant magnitude of inspiratory resistive load. The temporal onset of inspiratory resistive loading was coupled to the onset of inspiration via measured airflow from the spirometer and another controller box (labelled R). Figure adapted from Rieger et al., (2020) under a CC-BY license.

#### S7 Original German wordings in the RMCT

The Respiratory Metacognition of Control Task (RMCT) was conducted in German (English translation is provided in the main body of the manuscript). Below, we list the original German formulations of questions and response scale labels (visual analogue scales; VAS) underlying the behavioural readouts.

Trial-by-trial readouts:

- prediction ($\hat{y}_{\text{int}}$): “Schaffen Sie es, auf der Insel zu landen und damit einen Atemwiderstand zu verhindern?” VAS labels from “definitiv nicht (0%)” to “definitiv ja (100%)”.
- metacognitive judgement of perceived control over breathing resistances ($y_{\text{mc}}$): “Wie viel Kontrolle können Sie aktiv darüber ausüben, ob Sie einen Atemwiderstand erhalten?” VAS labels from “überhaupt keine Kontrolle” to “vollständige Kontrolle”.

Retrospective judgements that were queried after every 10th trial of the task:

- overall control over breathing ($y_{\text{c}}$): “Haben Sie das Gefühl, dass Sie in den vergangenen Durchgängen Kontrolle über Ihre Atmung hatten?” VAS labels from “definitive nicht” to “definitiv ja”.
- tolerance of inspiratory resistances ($y_{\text{tol}}$): “Wie leicht fiel es Ihnen, die erschwerte Atmung in den vergangenen Durchgängen zu tolerieren?” VAS labels from “überhaupt nicht leicht” to “sehr leicht”.

Immediately after the last trial of the task, participants were asked to rate

- the unpleasantness of breathing resistances during the task ($y_{\text{av}}$): “Wie unangenehm empfanden Sie die Atemwiderstände während der Aufgabe?” VAS labels from “überhaupt nicht unangenehm” to “maximal unangenehm”.
